# A common molecular mechanism for cognitive deficits and craving in alcoholism

**DOI:** 10.1101/2020.07.13.200519

**Authors:** Marcus W. Meinhardt, Simone Pfarr, Cathrin Rohleder, Valentina Vengeliene, Janet Barroso-Flores, Rebecca Hoffmann, Manuela L. Meinhardt, Elisabeth Paul, Anita C. Hansson, Georg Köhr, Nils Meier, Oliver von Bohlen und Halbach, Richard L. Bell, Heike Endepols, Bernd Neumaier, Kai Schönig, Dusan Bartsch, Rainer Spanagel, Wolfgang H. Sommer

## Abstract

Alcohol-dependent patients commonly show impairments in executive functions that facilitate craving and can lead to relapse. The medial prefrontal cortex, a key brain region for executive control, is prone to alcohol-induced neuroadaptations. However, the molecular mechanisms leading to executive dysfunction in alcoholism are poorly understood. Here using a bi-directional neuromodulation approach we demonstrate a causal link for reduced prefrontal mGluR2 function and both impaired executive control and alcohol craving. By neuron-specific prefrontal knockdown of mGluR2 in rats, we generated a phenotype of reduced cognitive flexibility and excessive alcohol-seeking. Conversely, restoring prefrontal mGluR2 levels in alcohol-dependent rats rescued these pathological behaviors. Also targeting mGluR2 pharmacologically reduced relapse behavior. Finally, we developed a FDG-PET biomarker to identify those individuals that respond to mGluR2-based interventions. In conclusion, we identified a common molecular pathological mechanism for both executive dysfunction and alcohol craving, and provide a personalized mGluR2-mechanism-based intervention strategy for medication development of alcoholism.

## Introduction

The consumption of alcoholic beverages is popular in many cultures, but is also an important cause of morbidity and mortality worldwide, accounting for 3 million deaths (5.3% of all deaths) annually *(1)*. While most people can control their drinking behavior, others become excessive drinkers and can eventually develop an alcohol use disorder (AUD). The defining features of AUD are a pattern of compulsive heavy alcohol use and a loss of control over alcohol intake. Clinical presentation of the disorder is highly heterogeneous, but is often accompanied by deficits in executive functions, i.e. impairments in higher cognitive abilities involved in self-control, regulation of emotions, motivation, working memory, decision making, attention, and cognitive flexibility *(2)*. Alcohol-induced deficits in executive functions usually persist into protracted abstinence and can therefore contribute to craving and subsequent relapse in abstinent patients *(2)*.

A brain region critically involved in cognitive flexibility and top-down (i.e. executive) control is the prefrontal cortex *(3–5)*. Clinically, excessive alcohol use causes damage in the prefrontal cortex, which is associated with enhanced craving for alcohol, as well as impaired cognitive functioning in humans *(6–10)*. Recently, group II metabotropic glutamate receptors have received growing attention in addiction research, due to their abundance in the pathway from the medial prefrontal cortex (mPFC) to the nucleus accumbens (NAc), which mediates drug craving and relapse *(11–13)* as well as cognitive flexibility *(14)*. The metabotropic glutamate receptor subtype 2 (mGluR2) is thought to be an especially important target of drug-induced neuroadaptations, as it has been reported that long-term exposure to drugs of abuse results in its downregulation and reduced function *(15, 16)*. Moreover, we have demonstrated that alcohol dependence, in both humans and rats, leads to a long-lasting reduction of mGluR2 expression, specifically in the infralimbic subregion of the mPFC, which is associated with a loss of control over alcohol-seeking behavior in rats *(8)*. In addition, it has also been shown that mGluR2 can modulate cognitive flexibility and its stimulation enhances cognitive flexibility in rats *(17)*. Therefore, we hypothesized that an mGluR2 dysfunction in the mPFC may comprise a common molecular mechanism for deficits in several behavioral domains.

To test this hypothesis, we assessed cognitive flexibility in alcohol-dependent rats using the attentional set shifting ( adapted from *20, 21)* and delay discounting tests *(20)*. As a rat model of alcohol addiction, we used an established model of alcohol dependence, in which rats receive chronic intermittent alcohol vapor exposure. This leads to intoxication levels similar to those seen in clinical alcohol addiction, and induces long-lasting behavioral as well as pronounced molecular changes in the brain. Most importantly, alcohol-dependent rats in this model show persistent escalation of alcohol self-administration, increased motivation to obtain alcohol, and increased relapse-like behavior *(21–23)*.

Here, we show that alcohol-dependent rats exhibit reduced cognitive flexibility, a deficit accompanied by reduced spine density and functional changes in glutamatergic neurons of the mPFC. Using a global mGluR2 deficient rat line *(24)*, a lentiviral mGluR2 overexpression approach, and a neuron-specific mGluR2 knockdown in the infralimbic cortex, we then demonstrate a critical role of mGluR2 for reduced cognitive flexibility in alcohol-dependent rats. In summary, we identify an mGluR2 deficit in the mPFC as a common pathological mechanism that is necessary and sufficient both for increased alcohol-seeking behavior and impaired cognitive flexibility in rats, identifying mGluR2 activation as a potential therapeutic mechanism in alcohol addiction. We next demonstrate in male and female rats the feasibility and efficiency of an mGluR2 medication development strategy for reducing relapse-like drinking using highly selective pharmacological tools *(25, 26)*. Given the large heterogeneity in treatment response to approved alcohol addiction pharmacotherapies *(27)*, we finally present a putative biomarker approach, that uses [^18^F]-fluorodeoxyglucose (FDG) positron emission tomography (PET) to identify responders to interventions targeting this mechanism.

## Materials and Methods

All experiments were done in accordance with the EU guidelines for the care and use of laboratory animals and were approved by the local animal care committee (Regierungspraesidium Karlsruhe, Karlsruhe, Germany).

### Animals

All animals were housed in groups of four under a 12 h light/dark cycle with food and water available *ad libitum* in home cages. The generation of CamKII-Cre transgenic rat, Grm2 *407 genotyping, and generation of a Cre-inducible mGluR2 knockdown virus is described in Supplementary Information.

### General experimental design

Six cohorts were used for behavioral studies. First, 18 CamKII-Cre transgenic rats were trained for operant alcohol-self-administration to test Cre-inducible shRNA-AAV mGluR2 knockdown in the infralimbic cortex. After reinstatement tests, rats underwent the ASST. Then four wistar batches (n = 32 each) were randomly assigned into two groups, which either underwent 8 weeks of chronic intermittent ethanol (CIE) exposure or air exposure. The first batch was used for the ASST, the second received bilateral Lenti-virus injections prior to the ASST, the third batch underwent the delay discounting task and the fourth batch was used for operant self-administration following mGluR2 agonist treatment. Finally, Indiana P (n = 12) and NP (n = 14) rats were directly tested on their ASST performance.

All other procedures, such as **behavioral procedures (**alcohol vapor exposure, attentional set shifting task (ASST), delay discounting test, operant alcohol self-administration, cue-induced reinstatement testing, and long-term voluntary alcohol consumption with repeated deprivation phases (ADE measurements)), **neuroanatomical experiments (i**mmunohistochemistry procedures, RNAscope fluorescent in-situ hybridization and spine density analysis), **biochemical Assays (**Dual-Luciferase assay, Western blot of mGluR2 and [^35^S] GTP-γ-S autoradiography), e**lectrophysiology**, magnetic resonance imaging (**MRI**) and positron emission tomography (**PET**) are described in Supplementary Information.

### Statistics

All data are expressed as mean ± SEM. Alpha level for significant effects was set to 0.05. Operant alcohol-seeking between-group data and ADE experiments were analyzed using repeated measures ANOVA, followed by Newman-Keuls post hoc test. Data of ASST subtasks were analyzed using one-way ANOVA. Overall group differences across all ASST subtasks were analyzed using repeated measures ANOVA. Spine density data were analyzed using two-tailed t-tests. Luciferase Assay and RNAscope in-situ hybridization data were analyzed using two-tailed t-tests. Western blot data were analyzed using one-tailed t-tests because of the expected reduction in protein levels in the mGluR2 knockdown group. For GTP-γ-S binding assays, values were calculated as percent of baseline value in the same region and animal and expressed as percent stimulation (± SEM) and were statistically analyzed by one-way ANOVA. For FDG-PET experiments two way ANOVAs were performed. To correct for multiple testing with 19,536 brain voxels, a threshold-free cluster enhancement (TFCE) procedure with subsequent permutation testing, thresholded at p < 0.05, was used *(28)* for all t-maps.

## Results

### Reduced cognitive flexibility after chronic intermittent alcohol exposure

Rats were made alcohol-dependent by chronic intermittent ethanol (CIE) vapor exposure for 7 weeks. During week 2 throughout 7, blood alcohol concentrations (BAC) were 250-300mg/dl. This exposure resulted in pronounced withdrawal signs compared to air-exposed animals (Figure 1A). After a prolonged abstinence period of 3 weeks the rats were used to assess executive functions. Sixteen alcohol-exposed and 16 air-exposed Wistar rats were tested in an attentional set shifting test (ASST) (Figure 1B), a rodent version of the Wisconsin card sorting task *(19, 29)*, a widely employed neuropsychological test in humans for assessing cognitive flexibility, with this deficit observed in alcohol-dependent patients *(30–32)*. Two alcohol-dependent and one control animal were excluded from the ASST, because they failed to learn the initial simple discrimination (SD) task. Repeated measures ANOVA revealed a main effect of alcohol exposure on alcohol-dependent and control animals’ ASST performance (F_[1,27]_ =11.34, p < 0.002) as well as a significant group x alcohol exposure interaction (F _[6,162]_ =2.62, p < 0.018). While there were no significant differences between the groups in SD, compound discrimination (CD), compound discrimination reversal (CDrev) and compound discrimination repetition (CDrep) tasks (one-way ANOVAs, SD: F_[1,26]_= 0.33, p = n.s.; CD: F_[1,26]_= 0.36, p = n.s.; CDrev: F_[1,26]_= 2,72, p = n.s.; and CDrep: F_[1,26]_= 0.001, p = n.s., respectively), we observed pronounced differences at higher cognitive demands. Thus, alcohol-dependent rats needed significantly more trials to reach criterion than non-dependent rats in the subtasks involving an intradimensional (IDS) or extradimensional shift (EDS) (one-way ANOVAs, IDS: F_[1,26]_= 5.99, p < 0.05; EDS: F_[1,26]_= 7.4, p < 0.05, respectively; Figure 1B). Thus, ASST performance was not generally impaired after CIE exposure but significant impairments were observed when the rules of the task were changed (IDS and EDS). This inability to adapt their strategies across testing conditions indicates a reduced cognitive flexibility in alcohol-dependent rats.

**Figure 1.**
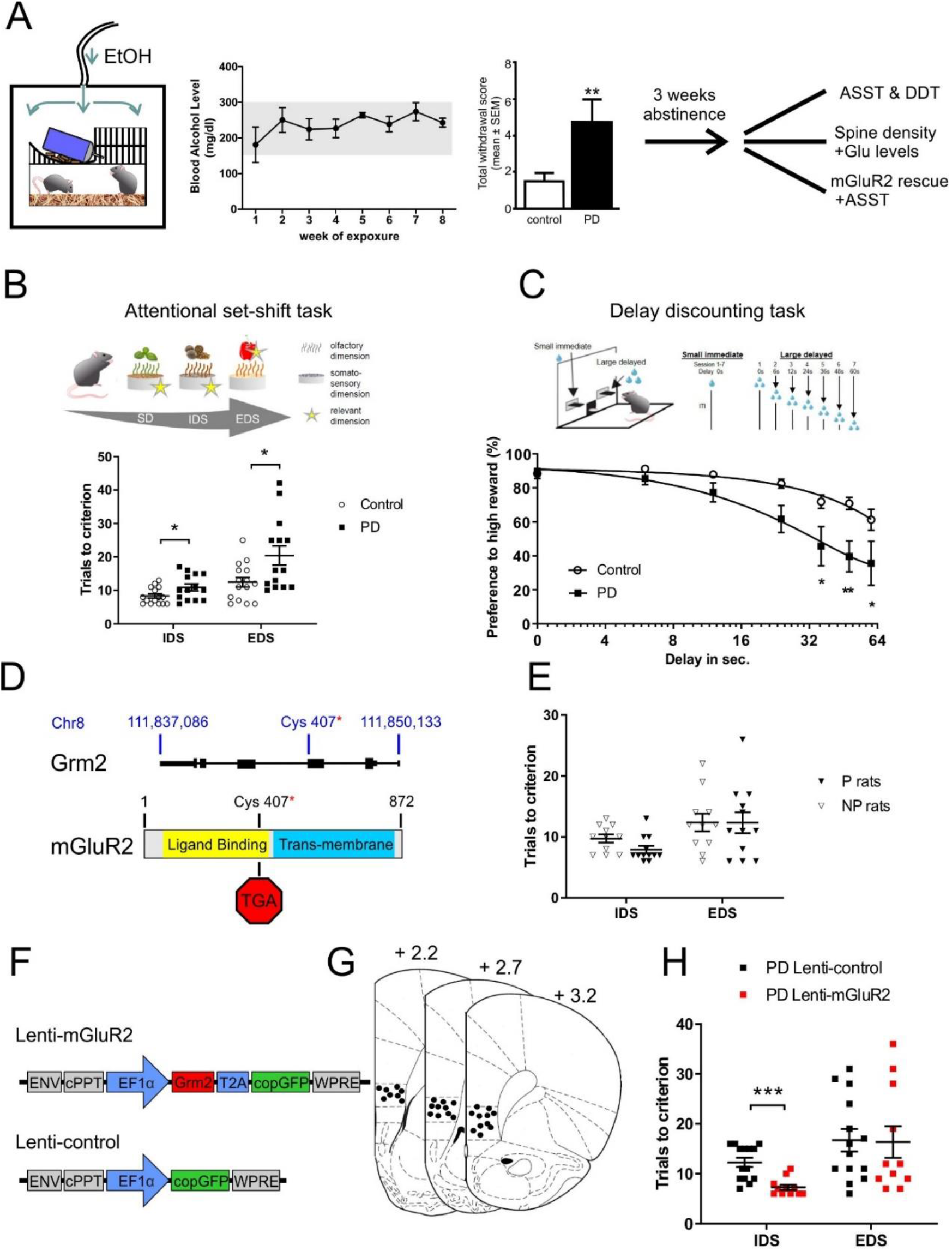
Alcohol-induced mGluR2 deficit within the infralimbic cortex leads to impaired ASST performance. **A)** Scheme of chronic alcohol vapor exposure procedure leading to blood alcohol concentrations (BAC) between 250 – 300mg/dl (for week 2-7) and significantly increased somatic withdrawal scores as compared to air-exposed control rats. After a prolonged abstinence period of 3 weeks three batches of animals were used either for spine density analysis, ASST or lentiviral mGluR2 rescue in the infralimbic cortex, followed by ASST. **B)** Scheme of ASST. The ASST setup consists of a start area and a choice area, containing two ceramic digging pots, separated by a central divider. The rats have to make serial discriminative choices based on olfactory (odors) or tactile (media) cues/ dimensions. The ASST paradigm consists of six sub-tasks divided into 4 test days characterized by increasing cognitive demands. The extradimensional set-shift is the most cognitive demanding sub-task, as the relevant dimension switches from tactile to olfactory. ASST performance of alcohol-dependent (post-dependent - PD) rats (n = 14, black squares) and control rats (n = 15, white circles). There was an overall significant difference between PD and control rats, with PD rats needing significantly more trials to criterion in the IDS and EDS subtasks. **C)** Scheme of the delay discounting task; one lever was randomly assigned as low reward lever (30μL/trial) and the second as high reward lever (90 μl/trial). The graph below represents the percentage preference for the high reward against the delay of the high reward in time. In comparison between the control and the alcohol-dependent rats, a steeper curve is observed with the dependent rats. Data are represented as mean ± SEM. **D)** Schematic representation of genomic location of premature stop-codon in Indiana P rats ( adapted from *24)*. **E)** There was no significant difference in the IDS/EDS tasks of ASST in alcohol-naïve Indiana P rats (n = 12, black triangles) compared to Indiana NP rats (n = 12, white triangles). **F)** Schematic representation of lentiviral constructs. Lenti-mGluR2 expresses Grm2 and copGFP under control of EF1α promoter. Lenti-control only expresses copGFP under EF1α control. **G)** Injection placements are represented by black circles. Injection sites were verified within the infralimbic cortex from +3.2 to +2.2mm anterior to bregma *(70)*. **H)** ASST performance of PD rats injected with either Lenti-mGluR2 (n = 16, red squares) or Lenti-control (n = 16, black squares) into the infralimbic cortex. In the EDS component of the task no difference was observed between control and virally infected PD rats. However, in the IDS component lenti-mGluR2 injected animals overall needed significantly less trials to criterion. IDS=intradimensional shift, EDS=extradimensional shift; *p < 0.05, ** p < 0.01, ***p < 0.001.

Reduced cognitive flexibility was also seen in a reward delay discounting test. Alcohol dependent individuals display greater discounting of future rewards than controls and this is accompanied by decreased activation of the executive system *(33–36)*. Paralleling clinical observations, the alcohol-dependent rats (n = 8/group) displayed a steeper discounting curve suggesting a faster switch in preference towards the lower over the delayed higher reward (repeated measurement ANOVA, group effect: F_[1,12]_= 7.78, p <0.05; delay effect: F_[6, 72]_= 29.72, p <0.001 and treatment x delay interaction: F_[6, 72]_= 3.84, p <0.01). The results of the ASST and delay discounting tests are consistent with a previous study in mice where CIE exposure led to reduced cognitive flexibility and the inability to alter behavioral responses under changing environmental demands *(7)*.

### Reduced cognitive flexibility is accompanied by structural and functional changes in the corticostriatal system

Given that CIE exposure induced structural neuronal changes within the PFC in mice *(7)* we next asked whether reduced cognitive flexibility is accompanied by structural and functional changes within the mPFC. We analyzed spine density in the mPFC of alcohol-dependent and air-exposed control rats and found significantly higher in the mPFC of alcohol-dependent rats compared to control rats (two-tailed t-test, t_[1,29]_ = 2.71, p < 0.011, Figure S1). Projections from the mPFC to the NAc mediate different aspects of the ASST *(14)*. In the NAc of alcohol-dependent rats we found a significantly lower spine density compared to control rats (t_[1,8]_ = 2.63, p < 0.03, Figure S1).

Next, we asked if these structural changes in the mPFC-NAc pathway were accompanied by functional alterations. It is known that glutamatergic neuroadaptations arise in corticostriatal projections after exposure to drugs of abuse including alcohol *(37)*. In particular, CIE exposure in mice leads to persistent changes in glutamatergic projection neurons from the mPFC *(7, 8)*. Given that infralimbic mPFC to NAc shell projections are in particular altered in alcohol-dependent rats and alcohol-dependent patients *(8)*, we measured baseline and alcohol-induced extracellular glutamate levels in the mPFC and NAc shell by microdialysis in freely moving rats (Figure S2A-F). In the mPFC, we found strongly reduced basal levels of glutamate (t_[1,169]_ = 10.65, p < 0.0001) indicative of a hypo-glutamatergic state. Systemic administration of increasing doses of alcohol (0, 0.5, 1.0, and 2.0g/kg i.p) led to robust and dose-dependent increases in extracellular glutamate levels in alcohol-dependent rats with no changes in non-dependent rats (repeated measures ANOVA time x treatment interaction: F_[18, 180]_= 2.02, p <0.01) (Figure S2A-C). In the NAc shell systemic administration of increasing doses of alcohol did not affect extracellular glutamate levels in alcohol-dependent or control rats (repeated ANOVA time x treatment interaction: F_[19,209]_= 1.08, n.s.) (Figure S2D-F). These findings are consistent with previously described persistent changes at glutamatergic projection neurons in the mPFC *(7, 8)* and further show a heightened sensitivity of the corticostriatal pathway to alcohol-stimulation. In summary, we found that alcohol-dependent rats exhibited persistent structural and functional alterations within the corticostriatal system. Whether these alterations, especially in the glutamatergic system, contribute to reduced cognitive flexibility in alcohol-dependent rats was studied in the next set of experiments.

### Improved ASST performance in alcohol-dependent rats after mGluR2 rescue in infralimbic cortex

Glutamatergic activity and homeostasis is regulated by metabotopic glutamate receptors 2 and 3 (mGluR2/3). In particular, these receptors typically provide negative feedback on glutamate actions: preferentially presynaptic mGluR2 inhibit glutamate release, while in glia, mostly mGluR3 increase glutamate uptake from the synapse by increasing the expression of glial glutamate transporters *(38)*. In our previous work we demonstrated a significant reduction of mGluR2 within the mPFC, especially in the infralimbic cortex in alcohol-dependent rats and alcohol-dependent patients *(8)*. This deficit in mGluR2 expression may explain the observed heightened sensitivity of glutamate release in response to alcohol in alcohol-dependent rats (Figure S2A, B). Here we studied the contribution of an mGluR2 deficit to reduced cognitive flexibility in the ASST. We first used a rat line that carries a spontaneously arising stop-codon introducing mutation in the in the Grm2 gene locus *(24* and Figure 1D) to study whether this global mGluR2 deficit would lead to similar impairments in ASST performance as chronic intermittent alcohol exposure. For this purpose we used 12 alcohol-naïve male Indiana alcohol preferring (P) rats and 14 male Indiana non-preferring (NP) rats. Two NP rats were excluded from the experiment, because they failed to learn the early stages of ASST. There was a significant difference on the first testing day during simple discrimination (F_[1,21]_ = 8.9, p < 0.007). However, there were no significant differences in the other subtasks of the ASST (CD: F_[1,21]_ = 1.49, n.s. ; CDrev: F_[1,21]_ = 0.07, n.s. ; CDrep: F_[1,21]_ = 0.74, n.s. ; IDS: F_[1,21]_ = 4.15, p = n.s. ; EDS: F_[1,21]_ = 0.0002, n.s.; Figure 1E). There was also no overall significant difference as confirmed by repeated measures ANOVA analysis (F_[1,21]_ = 1.55; n.s.). Thus, a global functional mGluR2 knockout does not appear to be associated with deficits in ASST performance likely due to compensatory mechanisms during development.

Next we asked whether a local rescue of mGluR2 expression and function within the infralimbic cortex in alcohol-dependent rats would lead to a normalization of cognitive flexibility. We applied a lentiviral overexpression (Figure 1F, G) approach in 32 alcohol-dependent Wistar rats. After CIE exposure, the animals received bilateral injections into the infralimbic mPFC with the respective lentivirus vectors (lenti-mGluR2 n = 16; lenti-control n = 16; seven rats excluded due to miss-targeting or minimal virus expression). The animals were allowed to recover for two weeks before the start of the ASST. As shown in Figure 1H, the impairments in alcohol-dependent rats in the IDS was reversed by mGluR2 rescue within the infralimbic cortex. Hence, there was a significant reduction in the IDS in the lenti-mGluR2 group, compared to lenti-control (F_[1,23]_ = 19.45, p < 0.0002). There were no significant differences in the other ASST subtasks (SD: F_[1,23]_ = n.s. ; CD: F_[1,23]_ = 0.14, n.s. ; CDrev: F_[1,23]_ = 1.35, n.s. ; CDrep: F_[1,23]_ = 0.06, n.s. ; IDS: F_[1,23]_ = 0.03, n.s. ; EDS: F_[1,23]_ = 0.008, n.s.). In summary, rescue of mGluR2 expression within the infralimbic cortex led to a rescue of cognitive flexibility in the IDS task.

### Reduced cognitive flexibility in alcohol-naïve rats after mGluR2 knockdown in the infralimbic cortex

The lentiviral overexpression approach shows that a rescue of ASST performance in alcohol-dependent rats is dependent on a rescue of mGluR2 expression and function within the infralimbic cortex. Our previous findings indicate that corticostriatal projection neurons to the NAc shell are particularily affected by chronic alcohol exposure and are characterized by an alcohol-induced mGluR2 deficit *(8)*. In the next set of experiments, we asked whether a deficit in mGluR2 function in infralimbic cortex projection neurons in alcohol-naïve animals is sufficient to mimic reduced flexibility as seen in alcohol-dependent rats. Very recently, transgenic rat models were introduced with neuron-specific genome modification in the adult rat brain *(39)*. We followed such an approach in order to specifically downregulate mGluR2 expression in projection neurons from the infralimbic cortex. For this purpose, we developed a novel CamKII-Cre transgenic rat line, expressing cre-recombinase under control of calcium/calmodulin-dependent protein kinase II alpha (CamKII) promoter (Figure 2A), which was demonstrated to drive expression predominantly in excitatory forebrain neurons *(40, 41)* in combination with a virally delivered shRNA targeted against mGluR2 mRNA (Figure 2D).

**Figure 2.**
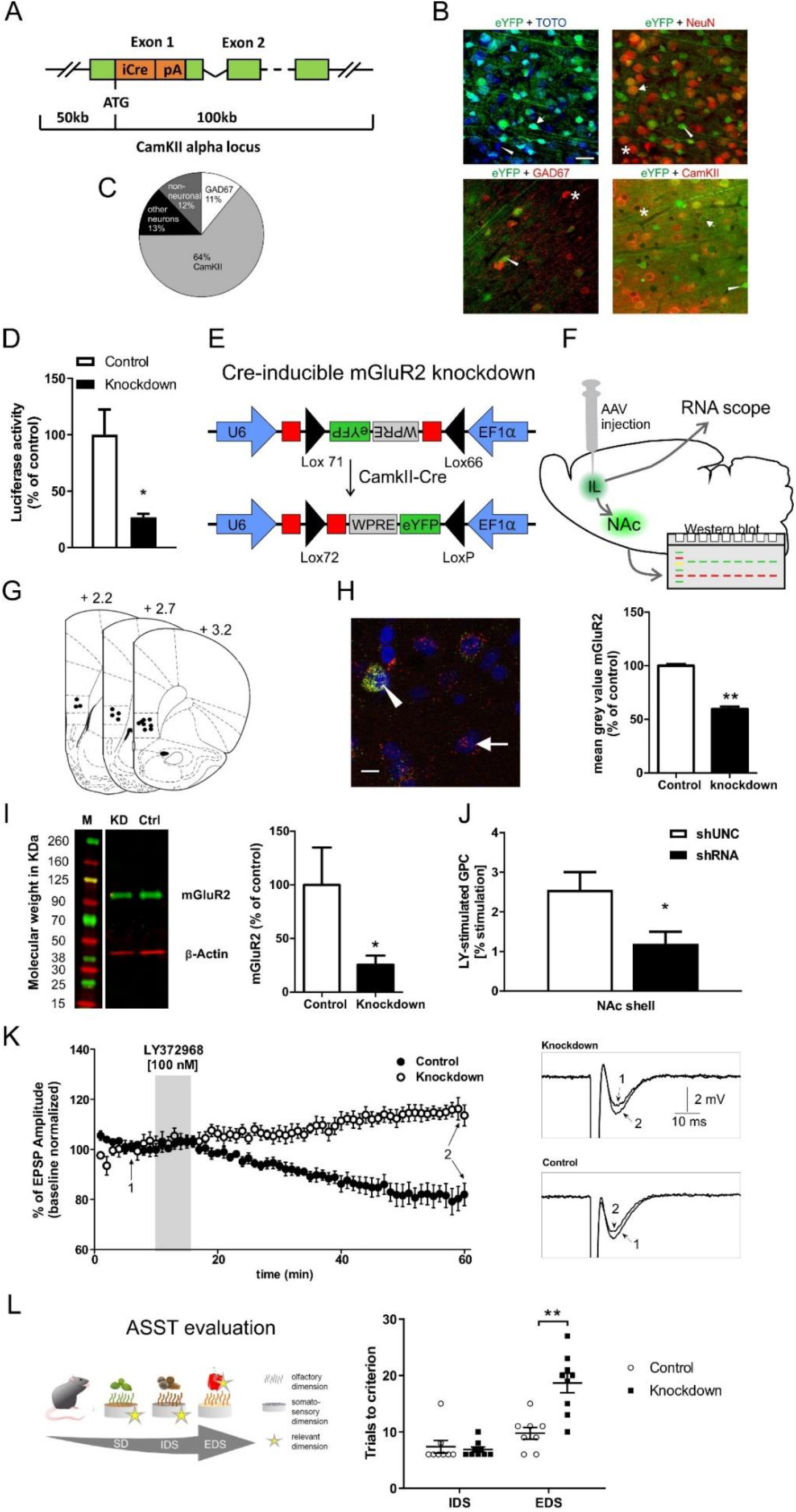
Characterization of specific mGluR2 knockdown in infralimbic cortex and corticostriatal projection neurons and impairments of executive functions in CamKII-mGluR2 knockdown rats. **A)** Bacterial artificial chromosome construct for neuronal specific Cre-expression in transgenic rats constitutively under control of the CamKII promoter. **B)** Representative images of eYFP co-localization with TOTO-3, NeuN, GAD67 and CamKII. Scale bar 20μm. Asterisks indicate single-positive cells for the respective cellular marker, triangles indicate single-positive cells for viral eYFP expression and arrows indicate double-positive cells for the respective cellular marker and the viral eYFP marker. **C)** Proportion of CamKII and GAD67 positive cells. **D)** Dual luciferase assay of control AAV (white bar) and Cre-inducible mGluR2 knockdown AAV (black bar). There was a significant downregulation of mGluR2 reporter luciferase activity when co-transfected with the knockdown AAV compared to control AAV. **E)** Schematic representation of the AAV construct. The shRNA coding sequence (in red) was split in the middle and inserted in opposite directions into the lox71 and lox66 flanked cassette. The reporter gene eYFP was also inverted and inserted opposite of the EF1α promoter. Without Cre-recombination no shRNA and no eYFP expression are possible. After Cre recombination shRNA expression is driven by U6 promoter and eYFP expression is driven by EF1α. **F)** Schematic representation of AAV injection into the infralimbic mPFC and projection to the nucleus accumbens (NAc). Knockdown of mGluR2 mRNA was detected via RNAscope in situ hybridization, protein levels were determined by western blot using NAc shell tissue punches (see I). **G)** Injection placements of Cre-inducible knockdown AAV and control AAV are represented by black circles. **H)** Left: Representative image of eYFP and mGluR2 fluorescent in-situ hybridization of CamKII-Cre rats injected with Cre-inducible mGluR2 knockdown AAV. Arrow indicates a single cell for mGluR2. Triangle indicates a double positive cell expressing eYFP and mGluR2. Scale bar: 10μm. Right: Quantification of Cre-inducible knockdown efficiency on mRNA level using fluorescent in-situ hybridization in CamkII-Cre rats. There was a significant reduction of mGluR2 mRNA transcripts in animals injected with the knockdown AAV (black bar) compared to control AAV (white bar). **I)** Left: Representative Western blot for mGluR2 and ß-actin in micropunched NAc shell tissue of CamKII-Cre rats after Cre-inducible knockdown AAV (KD) or control AAV (Ctrl) injection. M = Marker. Right: Quantification of mGluR2 protein levels in NAc shell of CamKII-Cre rats after injection of control (white bar) or Cre-inducible knockdown AAV (Black bar), by analyzing signal intensity. There was a significant downregulation of mGluR2 protein in infralimbic cortex after knockdown AAV injection. *p < 0.05; **p < 0.01. **J)** There was decreased LY379268-stimulated G-protein coupling (GPC) in the NAc shell following infralimbic cortex mGluR2 knockdown, compared to controls (n = 4/group; shUNC baseline value: 291±20 nCi/g, shRNA baseline value: 279±9 nCi/g, mean±SEM). **K)** Normalized EPSP amplitude (mean +/- SD) before (baseline), during the 5 min application of 100 nM of the mGluR2 agonist LY379268 (gray shadow) and during prolonged LY379268 washout. Representative voltage traces of control and knockdown are shown (“1” labels baseline traces; “2” labels traces after 45 min LY379268 washout). **L)** ASST performance of CamKII-Cre rats injected with either control AAV (n = 9, white bars) or Cre-dependent knockdown AAV (n = 9, black bars) into the infralimbic mPFC. The knockdown animals needed significantly more trials to criterion in the EDS subtask and overall needed more trials to criterion as compared to control animals. **p < 0.01.

First we characterized the Cre expression pattern of CamKII-Cre rats in the infralimbic cortex by bilateral injections of a Cre-inducible AAV expressing the fluorescent marker eYFP. Four weeks after the AAV injection rats were killed and brain sections were stained with TOTO-3 for nuclear counterstaining. The cellular location of the fluorescence marker was analyzed by immunolabeling for the neuronal marker NeuN, the interneuron marker GAD67, and CamKII as a marker for prefrontal projection neurons. We found that ~43% of all TOTO-3 cells colocalized with eYFP expression, whereas 52% of all neurons (NeuN) colocalized with eYFP. Of all eYFP positive cells, ~88% were NeuN positive, ~64% were CamKII positive and ~11% were GAD67 positive (Figure 2B, C). Thus, there is no exclusive co-localization of Cre with CamKII neurons in the infralimbic cortex of CamKII-Cre rats. However, the majority of Cre-expressing neurons is CamKII positive, which indicates this transgenic rat line provides a strong preference for manipulations of projection neurons from the infralimbic mPFC.

We then tested the knockdown efficiency of a newly designed conditional mGluR2 shRNA expression system in-vitro and in-vivo on the mRNA and protein levels, as well as by GTP-γ-S autoradiography and local field potential (LFP) recordings (Figure 2D-K). For initial verification of the knockdown strategy, a dual luciferase assay was performed with the in-vitro recombined Cre-dependent shRNA AAV expression vector, together with a firefly luciferase reporter construct, harboring the mGluR2 shRNA target sequences. The mGluR2 knockdown-AAV construct reduced firefly luciferase activity to ~27% compared to control AAV (t_[1,4]_ = 3.255, p < 0.03; Figure 2D). We then tested the knockdown efficiency of the Cre-inducible knockdown AAV in-vivo, CamKII-Cre rats were injected with the mGluR2 shRNA-AAV or control AAV (Figure 2E-G). After four weeks of virus expression mRNA levels of mGluR2 were analyzed using fluorescent in-situ hybridization. Compared to the control AAV, the shRNA expressing AAV significantly reduced mGluR2 mRNA levels by ~40% (t_[1,2]_ = 16.8, p < 0.004; Figure 2H). Knockdown efficiency of the Cre-dependent knockdown AAV on the protein levels was tested by Western blots of NAc shell tissue samples (n = 8/group). Compared to control AAV, the knockdown AAV significantly reduced mGluR2 protein levels by ~75% (t_[1,14]_ = 1.84, p < 0.04; Figure 2I). We also tested the effect of the mGluR2 knockdown on downstream signaling. G-protein activation by the mGluR2 agonist LY379268 was measured by [^35^S]GTP-γ-S autoradiography (Figure 2J). Compared to the control shRNA, infralimbic cortex injections of mGluR2 knockdown shRNA significantly reduced mGluR2 receptor binding in the NAc shell (One-way ANOVA: F_[1,6]_ = 6.28, p < 0.05). Finally, we tested the mGluR2 knockdown at a functional level by recording field excitatory postsynaptic potentials (EPSPs) in slice preparations containing NAc shell and infralimbic cortex. Pre-synaptically located mGluR2 act as auto-receptors *(42)* and their activation induces long-term depression (LTD) of glutamatergic transmission *(24, 43)*. Here, we activated mGluR2 pharmacologically using 100 nM of the mGluR2/3 agonist LY379268 (Figure 2K). EPSPs were recorded in NAc shell by stimulating glutamatergic neurons in the infralimbic cortex. Bath perfusion of LY379268 (5 min) reduced the amplitude of EPSPs in control rats by ~20% but increased the amplitude of EPSPs in knockdown rats by ~10 %, suggesting an uncompensated loss of mGluR2 function (Figure 2K). In summary, our site-specific mGluR2 receptor knockdown could be successfully demonstrated on various levels - in-vitro, in vivo as well as pharmacologically and functionally. We conclude that, following our targeting strategy, infralimbic corticostriatal projection neurons showed the pronounced deficit in mGluR2 function similar to that observed in alcohol-dependent rats.

To test if a deficit in mGluR2 function in infralimbic cortex and its corticostriatal projection neurons affects ASST performance and would mimic an alcohol-dependent phenotype, we injected CamKII-Cre rats (n = 18) with the knockdown AAV or the control AAV. An overall significant difference between the control and knockdown group was seen as confirmed by a repeated measures ANOVA (F_[1,15]_ = 9.02, p < 0.009) (Figure 2L). There were no significant differences in ASST subtasks SD – IDS as confirmed by one-way ANOVA (SD: F_[1,15]_ = 2.85, p = n.s. ; CD: F_[1,15]_ = 0.16, n.s. ; CDrev: F_[1,15]_ = 0.16, n.s. ; CDrep: F_[1,15]_ = 1.18, n.s. ; IDS: F_[1,15]_ = 2.78, n.s. . However, there was a significant difference between the groups in EDS performance (F_[1,15]_ = 18.07, p < 0.001; Figure 2L). Thus, as in alcohol-dependent rats (Figure 1C), there was a pronounced increase in EDS performance trial numbers indicative of reduced cognitive flexibility. These experiments demonstrate that a knockdown of mGluR2 in infralimbic corticostriatal projection neurons is sufficient to reduce cognitive flexibility in alcohol-naïve rats in a manner that mimics that found in the alcohol-dependent rats.

### Cre-dependent mGluR2 knockdown in the infralimbic cortex is sufficient to induce excessive alcohol seeking in non-dependent rats

In our previous study, we found that excessive alcohol-seeking behavior in rats with a history of alcohol dependence could be normalized by re-expressing mGluR2 in the infralimbic cortex *(8)*, demonstrating that reduced mGluR2 levels within the infralimbic cortex are necessary to induce this phenotype. To test sufficiency, we here used the above described viral mGluR2 knockdown strategy to reduce infralimbic cortex mGluR2 levels in non-dependent animals. We trained 19 CamKII-Cre transgenic rats to self-administer a 10% alcohol solution. After reaching a stable self-administration baseline (BL), all animals underwent extinction training (EXT) and one cue-induced reinstatement session before AAV injection (RE1, Figure 3A). Animals were ranked based on their RE1 performance and divided into two groups with matched performance. There were no significant differences between the prospective experimental groups during BL, EXT and RE1 (Figure 3B, Supplementary Table 1).

**Figure 3.**
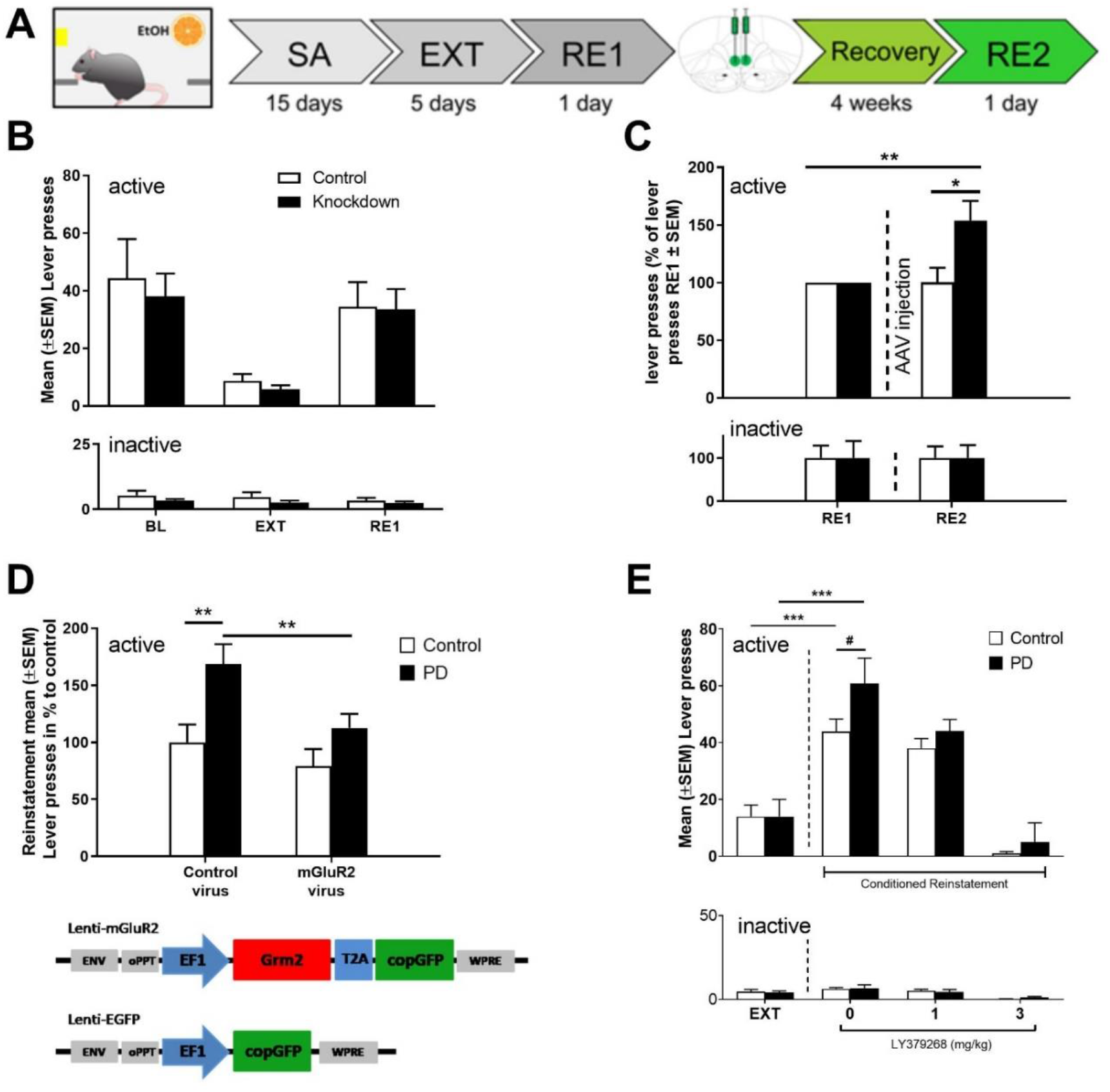
Effect of mGluR2 knockdown in infralimbic mPFC and corticostriatal projection neurons on cue-induced reinstatement of alcohol-seeking-behavior and a comparison with the alcohol-dependent phenotyp. **A)** Experimental timeline: all animals underwent alcohol self-administration (SA) and extinction (EXT) training, followed by a cue-induced reinstatement session of alcohol-seeking (RE1). Following stereotaxic AAV injection (control AAV or mGluR2 knockdown AAV, n = 9/group) and a recovery period of 4 weeks the animals were tested during a second cue-induced reinstatement session (RE2). **B)** Active and inactive operant responses of CamKII-Cre rats before control (white bars, n = 7) and knockdown (black bars, n = 7) AAV injection (two control and two knockdown animals had to be excluded from analysis, because they failed to reach the criterion for successful cue-induced reinstatement of alcohol-seeking, which was >10 active lever presses). There was no significant difference between the groups during baseline (BL), extinction (EXT) and cue-induced reinstatement (RE1) responding. **C)** Performance of CamKII-Cre rats injected with control or Cre-induced mGluR2 knockdown AAV. Data were normalized to RE1 performance (before AAV injection). There was a significant difference between the groups in RE2 and a significant difference in the knockdown group between RE1 (before AAV injection) and RE2. **D)** *Upper*: Lenti-mGluR2 significantly attenuated drug-seeking behavior only in alcohol-dependent rats (post-dependent – PD) down to the level of the control group. *Lower*: Schematic representation of the lentiviral expression plasmids used for the production of Lenti–mGluR2 and Lenti-EGFP. Presentation of the CS+ elicited significant reinstatement in both control and alcohol-dependent rats with lenti-EGFP ( Data are adapted from *9)*. **E)** Cue-induced reinstatement of alcohol-seeking behavior was tested after systemic administration of either vehicle or 1, and 3mg/kg i.p. of the mGluR2/3 agonist LY379268, which produced a dose-dependent decrease in alcohol-seeking behavior, especially in alcohol-dependent rats. *p < 0.05, **p < 0.01, *** p < 0.001 indicate significant differences between control, knockdown / alcohol-dependent (PD) rats and between RE2 and RE1, respectively.

Next, rats were bilaterally injected with control or Cre-dependent mGluR2 knockdown AAV (n = 9/group; four rats had to be excluded post-hoc due to incorrect placement or minimal virus expression). Four weeks after AAV injection, the animals were again tested for their operant alcohol-seeking performance. There was a significant difference within the knockdown group between RE1 and RE2, i.e. before and after AAV injection, respectively (t _[1,11]_ = 3.2, p < 0.01). For comparison of the effect size, we re-analyzed our previous experiment, that was done in a cohort of alcohol-dependent and control rats (n = 15/group) injected with an mGluR2 overexpression or control lenti-vector, which were tested for cue-induced reinstatement several weeks after cessation of CIE exposure *(8)*. As shown in Figure 3D, relative changes between alcohol-dependent and control rats were of similar magnitude as that for the infralimbic mPFC mGluR2 knockdown rats used here, and this effect was fully reversed by re-expression of mGluR2 in the infralimbic cortex of those alcohol-dependent rats. Thus, we demonstrate a bidirectional modulation of drug-seeking behavior – infralimbic cortex mGluR2 knockdown increased alcohol-seeking, while overexpression in mGluR2 deficient alcohol-dependent rats reduces it. From this direct comparison, we conclude that a downregulation of mGluR2 in infralimbic corticostriatal projection neurons is necessary and sufficient to induce an alcohol-dependent phenotype with respect to excessive alcohol-seeking behavior.

Taken together, we have identified a common pathological mechanism for two co-occurring phenotypes in addiction, excessive alcohol seeking and impaired cognitive flexibility. These results identify mGluR2 function in the mPFC as a target for medication development for alcoholism. Given that during withdrawal and protracted abstinence reduced executive function and craving can result in relapse, we next studied various systemic pharmacological interventions targeting the described mGluR2 deficit.

### mGluR2 mechanism-based interventions to treat alcohol relapse

We first targeted excessive alcohol-seeking in alcohol-dependent rats with the non-selective mGluR2/3 agonist LY379268. We trained 8 alcohol-dependent and 8 control rats to self-administer a 10% ethanol solution over 6 weeks (Figure S3). As described above, after stable responding and extinction all rats underwent cue-induced reinstatement testing. During the extinction phase, all rats reduced their active lever pressing to 10% or less of the previous baseline in a similar manner. Cue-induced reinstatement of alcohol-seeking behavior was then tested after systemic administration of either vehicle, 1 or 3mg/kg i.p. LY379268. The drug was injected 30 min before the start of the session. LY379268 produced a dose-dependent decrease in alcohol-seeking behavior, especially in alcohol-dependent rats (factor dose: F_[3, 42]_ = 49,01, p < 0.0001). The 3mg/kg dose produced sedative effects, as shown by the fact that most of the animals completely stopped responding and inactive lever responses were also affected (Figure 3E). This finding is in line with a previous report *(44)*. These authors reported enhanced attenuation of cue-induced reinstatement in alcohol-dependent rats by the mGluR2/3 agonist LY379268 within the same dose range as used here.

In the subsequent experiments we also used another established rat model of relapse - the alcohol deprivation effect (ADE) model - to test if the mGluR2/3 agonist LY379268 would also affect relapse-like drinking (Figure 4A). Alcohol deprivation is one factor that is known to considerably affect voluntary alcohol intake. Renewed access to alcohol solutions after a period of deprivation for several days/weeks leads to a pronounced, although temporary, increase in voluntary alcohol intake in rats - the so-called ADE *(45)*. Following repeated deprivation phases, the ADE is characterized by an increased motivation for and excessive pursuit of alcohol and therefore resembles a typical lapse or relapse situation in alcohol-dependent patients *(45, 46)*. This model has been widely used in examining the efficacy of pharmacological agents in preventing alcohol relapse *(46)*.

**Figure 4.**
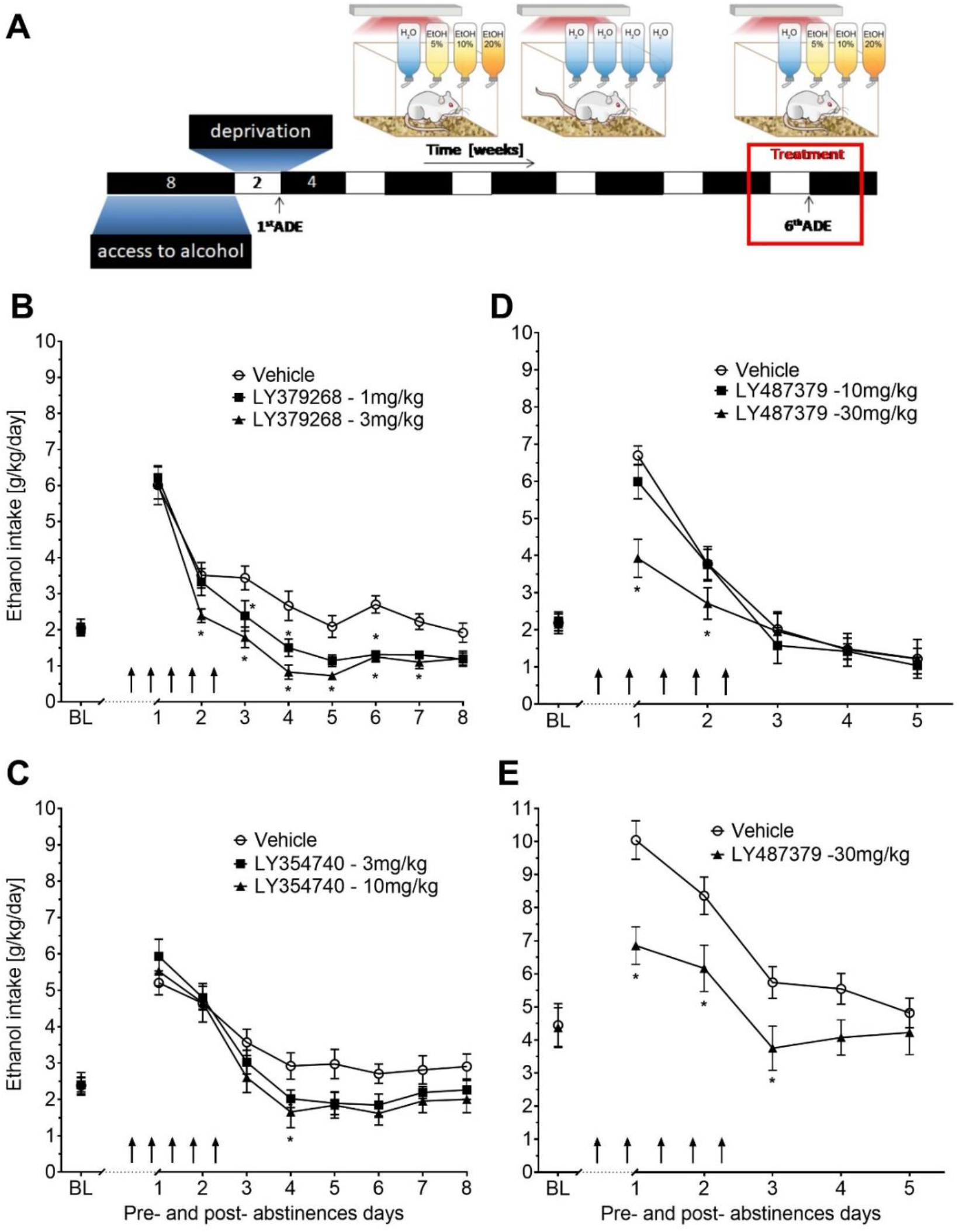
Effects of mGluR2/3 agonists and an mGluR2 PAM in an animal model of relapse. **A)** Experimental timeline – all animals underwent repeated cycles of alcohol consumption (in a free choice paradigm with water, 5%, 10%, or 20% ethanol solutions) and deprivation phases (only water available). After the 6^th^ alcohol deprivation phase pharmacological intervention experiments were conducted. **B-E)** Total alcohol intake (g/kg/day) before and after an alcohol deprivation period of three weeks in the ADE model. Arrows indicate the administration of **B)** vehicle (n = 8), 1 mg/kg of LY-379268 (n = 8) or 3 mg/kg of LY-379268 (n = 8), **C)** vehicle (n = 10), 3 mg/kg of LY-354740 (n = 8) or 10 mg/kg of LY-354740 (n = 7) in male rats and administration of either vehicle or LY487379 in both (**D**) male and (**E**) female rats [vehicle (n = 8 males, n = 10 females), or 10 mg/kg of LY487379 (n = 7 males), or 30 mg/kg of LY487379 (n = 7 males, n = 8 females)]. The last three days average of alcohol intake are given as baseline drinking – “BL”. Data are presented as means ± S.E.M. * indicates significant differences from the vehicle group, p < 0.05.

Following the re-introduction of alcohol solutions after a period of abstinence, the vehicle-treated group showed a typical increase in alcohol consumption, indicating the occurrence of an ADE. With respect to the LY379268 treatment, a two-way ANOVA for repeated measures revealed a significant increase in alcohol intake after a deprivation phase in all animal groups as compared to basal drinking (factor day: F_[8,168]_=121.5, p < 0.0001) (Figure 4B). The LY379268 treatment significantly reduced the expression of the ADE (factor day × treatment group: F_[16,168]_=2.7, p < 0.001). No significant difference in water intake (Figure S4A) was seen. LY379268 treatment did not lead to significant changes in body weight, showing that food intake or metabolism was not altered during the treatment days (Supplementary Table 2). Locomotor activity data were analyzed using recordings of 12-hour post-injection intervals that corresponded animals’ active phase. Overall, there was a general reduction in home-cage activity seen in all animal groups following re-gained access to alcohol. However, two-way ANOVA did not reveal significant changes in activity of LY379268 treated animals, when compared to the vehicle-treated rats (factor treatment group: p = n.s.; and factor day × treatment group: p = n.s.; Figure S5A). These data together with the recordings of the animal’s body weight suggests that repeated administration of LY379268 can lead to nonspecific treatment effects – resulting in sedation – at a high dose *(47)*.

Next we tested LY354740, a structural analog of LY379 with an six times higher ability to discriminate between mGluR2 and mGluR3 *(48)*. The LY354740 treatment also significantly reduced expression of the ADE [factor day × treatment group: F_[16,176]_=2.1, p < 0.01] (Figure 4C). No significant difference in water intake (Figure S4B) was seen. LY354740 treatment did not lead to significant changes in body weight, showing that food intake or metabolism was not altered during the treatment days (Supplementary Table 3). Overall, there was a general reduction in home-cage activity in all animal groups following re-gained access to alcohol. However, two-way ANOVA did not show any significant changes in activity of LY354740 treated animals, when compared to the vehicle-treated rats [factor treatment group: p = 0.3; and factor day × treatment group: p=0.7] (Fig. S5B). Similar to the results with LY379268 treatment these data show that repeated administration of LY354740 leads to occurrence of minimal, if any, nonspecific treatment effects.

Anti-relapse treatment with an mGluR2/3 agonist would most likely require long-term treatment but it has been reported that chronic administration of group II mGlu receptor agonists can induce robust tolerance *(49)*. Therefore, subsequent ADE measurements were performed to test for the development of tolerance and for studying persistent treatment effects in a drug-free period. For this purpose, two groups of rats that were either treated repeatedly with vehicle or LY354740 were studied. Measurement of weekly alcohol intake showed that alcohol consumption was significantly different over the entire time course of the experiment (factor week F_[14,182]_=4.5, p < 0.0001; Figure S6). Subsequent post-hoc analysis revealed that LY354740 treatment reduced alcohol consumption during all three post-abstinence weeks and subsequent baseline drinking tended to be lower compared with the vehicle-treated group [factor treatment group: p=0.2 and treatment group × week interaction effect: p=0.3] (Figure S6). These data show that repeated sub-chronic treatment with an mGluR2/3 agonist does not induce tolerance with respect to its anti-relapse properties.

Although the results with mGluR2/3 agonists on relapse drinking are promising, LY379268 and other orthosteric group II mGluR agonists do not readily discriminate between contributions of mGluR2 and mGluR3 binding and seem to have long-term toxicity. Thus, it is unlikely that orthosteric agonists of these receptors can be developed for clinical use. To overcome these limitations, positive allosteric modulators (PAMs) selective for mGluR2 were recently developed *(26, 49)*. These PAMs exert their effects on mGluR2s through a binding site that is topographically distinct from that bound by the endogenous ligand *(50)* and potentiate receptor activation in the presence of glutamate *(51)*. LY487379 is a highly specific PAM that potentiates the effect of glutamate on the mGluR2 with little effect on other mGluR subtypes *(52)*. We tested this PAM in the ADE model and also evaluated the potential for sexually dimorphic effects, since sex differences are often seen in the efficacy of alcoholism treatment *(53)*. A two-way repeated measures ANOVA showed a significant increase in alcohol intake after a deprivation phase in all animal groups as compared to basal drinking [factor day: F_[5,95]_=98.2, p < 0.0001 and F_[5,80]_=45.0, p < 0.0001, for male and female rats, respectively] (Figure 4D-E). LY487379 treatment significantly reduced expression of the ADE in both male and female rats [factor day × treatment group: F_[10,95]_=4.5, p < 0.0001 and F_[5,80]_=4.9, p < 0.001, for male and female rats, respectively]. No significant difference in water intake (Figure S4) was seen, neither in male nor female rats. LY487379 treatment did not lead to significant changes in the body weight in male rats, showing that food intake or metabolism was not altered during the treatment days (Supplementary Table 4). However, the administration of LY487379 significantly reduced the body weight of female rats by nearly 2% [factor treatment group: t_[1,16]_=2.9, p < 0.05]. Overall, there was a general reduction in home-cage activity seen in all animal groups following re-gained access to alcohol. However, two-way ANOVA did not show significant changes in activity of LY487379 treated male animals when compared to the vehicle-treated rats [factor treatment group: p = 0.5; and factor day × treatment group: p=0.6] (Figure S5). These data, together with the recordings of the animals’ body weight, suggest that repeated administration of LY487379 leads to minimal, if any, nonspecific effects, such as sedation. Together with other data showing that PAMs can reduce alcohol-seeking behavior *(54)* we conclude that mGluR2 PAMs should be considered for clinical trials in alcohol-dependent patients.

### Development of a candidate biomarker for personalized treatment of individuals with an mGluR2 deficit

A challenge for medication development in alcoholism is the heterogeneity of the clinical population. The heterogeneity makes it unlikely that targeting a single pathophysiological mechanism among the multitude of those known to be involved in alcohol addiction, will be uniformly effective *(55)*, will be beneficial for all patients. A predictive biomarker able to identify patients responsive to interventions targeting mGluR2 would be a major advance, and improve the prospects of successful clinical development. In the next set of experiments, we propose an FDG-PET approach that may offer a biomarker strategy (Figure 5A).

**Figure 5.**
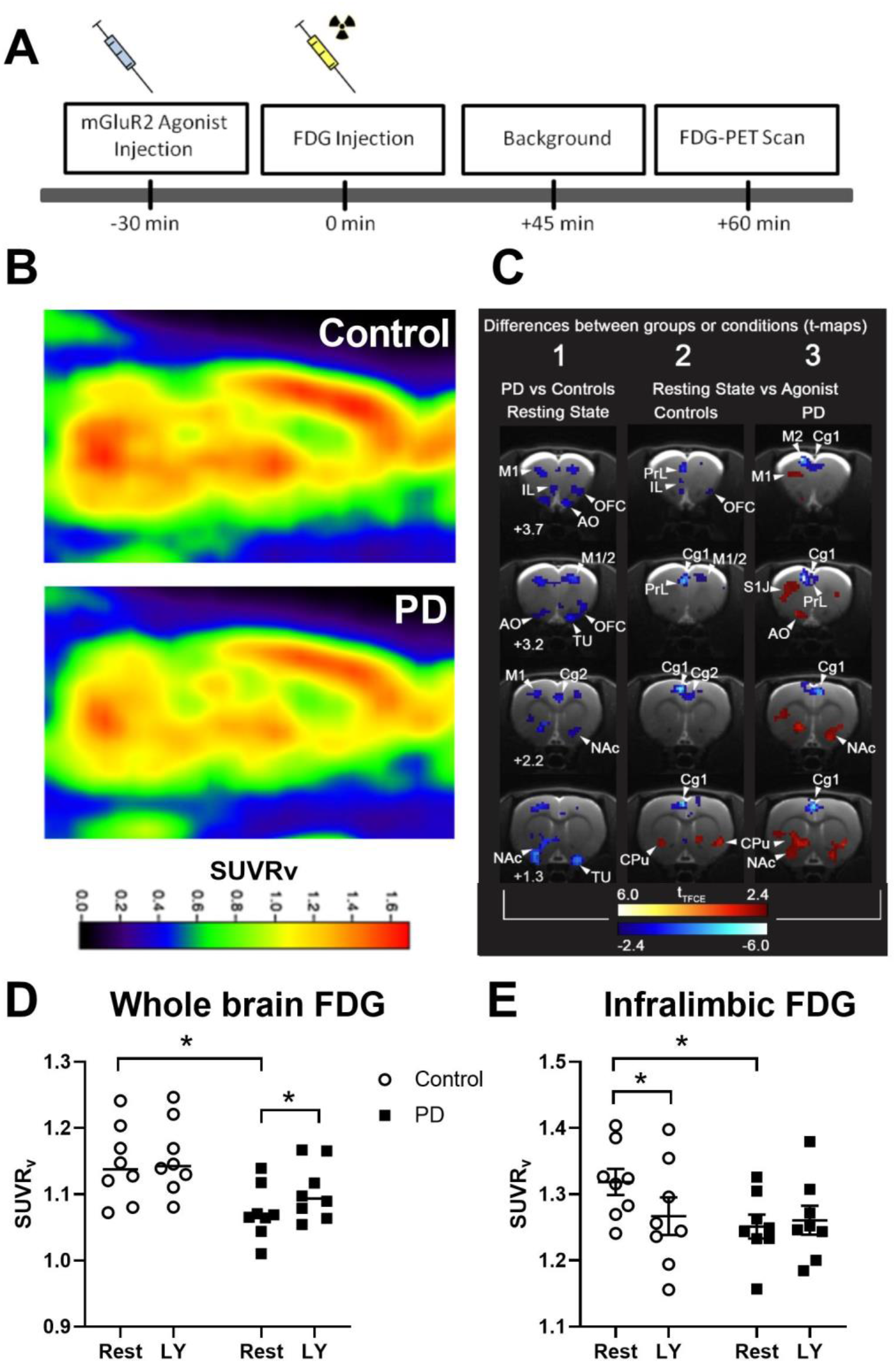
[^18^F]-FDG-PET in alcohol-dependent and control rats after application of the mGluR2/3 agonist LY379268. **A)** Experimental timeline: Animals received either vehicle or the mGluR2/3 agonist LY379268 (n = 8 per group) 30 min before [^18^F]-FDG injection. After the injection of [^18^F]-FDG, rats were placed in their home cage for 45 min followed by a PET scan. **B**) Representative normalized PET images of control and alcohol-dependent (PD) rats in a sagittal section. **C**) Differences between groups (PD vs. Control) or conditions (Resting condition or mGluR2/3 agonist treatment) represented in t-maps. White numbers indicate bregma levels. tTFCE bars are ranging from deep red to light red indicating significant increases in [^18^F]-FDG uptake ratio whereas deep blue to light blue indicates significant decreases. **D**) Whole-brain analysis between PD and controls shows a significant decrease in [^18^F]-FDG uptake (given as standardized uptake value ratio, SUVRv) in PD rats. **E**) mGluR2/3 agonist LY379268 (LY) challenge led to a decrease in [^18^F]-FDG uptake ratio (LY vs. vehicle) under resting state (rest) conditions in the infralimbic cortex of control rats, while no effect of the drug was found in the alcohol-dependent group. Cg1: cingulate cortex area 1, M1: primary motor cortex, M2: secondary motor cortex, PrL: prelimbic cortex, IL: infralimbic cortex, NAc: nucleus accumbens, OFC, orbitofrontal cortex, AO: accessory olfactory bulb, CPU: caudate putamen, TU: olfactory tubercle, S1J: primary somatosensory cortex. Data are presented as means ± S.E.M. * indicates significant differences from the vehicle group, p < 0.05.

The positron emission tomography (PET) radiotracer [^18^F]-fluorodeoxyglucose (FDG) is widely used in clinical diagnostics of CNS tumors, but also offers a valuable tool with which to identify impaired brain glucose metabolism in psychiatric conditions such as addiction *(56)*. Indeed several [^18^F]-FDG PET studies in alcohol-dependent patients have consistently reported decreases in overall brain glucose metabolism and FDG-PET has been proposed for biomarker development *(27)*. We have used FDG-PET to compare brain glucose uptake in alcohol-dependent rats, which have an mGluR2 deficit in the infralimbic cortex, vs. control non-dependent rats. As described in AUD patients, we found a reduction in glucose utilization in the whole brain of alcohol-dependent rats compared to controls (factor group: F_[1,14]_=6.4, p<0.05; Figure 5B-D). This effect could be reversed by LY379268 treatment (factor treatment: F_[1,14]_= 21.8, p<0.001).

In a refined region of interest (ROI) analysis, we focused on the mGluR2-impaired infralimbic cortex. In control rats the regional FDG signal was decreased by LY379268. In contrast, no such effect of LY379268 was found in the infralimbic of alcohol-dependent rats (Figure 5E). A two-way ANOVA revealed a significant treatment effect (F_[1,14]_=2.56, p < 0.01) as well as significant treatment x group interaction (F_[1,14]_=5.4, p < 0.05). Post-hoc analysis demonstrated a significant difference between control and alcohol-dependent rats under baseline condition (p < 0.05), confirming the results observed for whole brain analysis, as well as a significant difference between baseline and agonist treatment only in controls (p < 0.01). We conclude that alcohol-dependent rats with an mGluR2 deficit have blunted FDG response to the LY379268 challenge within the infralimbic cortex. Thus, the here proposed biomarker strategy should be able to identify patients with a regional decreased sensitivity to mGlu2 agonism, which are likely to respond to pharmacological treatment with a PAM to improve executive functions, alcohol craving and ultimately reduced relapse rates.

## Discussion

In this work, we use a translational multi-level approach to characterize the role of mGluR2 receptors in alcohol addiction. Using advanced genetic tools and pharmacologically validated rat models of AUD we first provide evidence that an mGluR2 deficit in the medial prefrontal cortex is both necessary and sufficient for diminished cognitive flexibility and increased drug craving. These findings establish a common molecular-pathological mechanism for two, mostly co-occurring intermediate phenotypes in addicted subjects. Prefrontal mGluR2 deficits can be caused by excessive alcohol as well as cocaine exposure *(8, 16, 57)*. We then demonstrate that pharmacological targeting of mGluR2 dysfunction may provide an effective strategy for relapse prevention in AUD. Given the large heterogeneity of AUD patient populations, successful pharmacotherapy have to be based on personalized medicine approaches. To identify potential treatment responders, i.e. those subjects that have an mGluR2 deficit, we propose a PET-based biomarker strategy and demonstrate its feasibility.

In the first set of experiments we examined executive functions, in particular cognitive flexibility of alcohol-dependent rats. In the ASST as well as in the delay discounting test we showed reduced cognitive flexibility in alcohol-dependent rats. This is in line with findings in alcohol-dependent patients who discount temporally distant rewards more than controls, a cognitive deficit that is accompanied by a decreased activation of the executive system ( i.e. mPFC; *34, 35)*. In summary, we show reduced cognitive flexibility in alcohol-dependent rats – a finding that is consistent with studies in alcohol-dependent mice *(7)* and patients.

By using genetic manipulations, including a rat model with an infralimbic cortex specific knockdown of mGluR2, we were able to modulate cognitive flexibility bi-directionally. These findings are consistent with improved ASST performance in rats treated with the mGluR2 positive allosteric modulator LY487379 *(17)*. In addition, our novel transgenic rat model expressing Cre-recombinase constitutively under CamKII promoter control provided a valuable addition to the recently published tool box of transgenic rats supporting neuron-specific genome modification in the adult rat *(39)*. In conclusion, an mGluR2 deficit is necessary and sufficient to mediate reduced cognitive flexibility in alcohol-dependent rats.

Using the same genetic manipulations, we further demonstrated that an mGluR2 deficit is also necessary and sufficient for enhanced cue-induced alcohol-seeking in rats. Alcohol-seeking is a critical component of alcohol craving *(46)* and thus we suggest that an mGluR2 deficit may also be involved in cue-elicited craving responses in humans. Reduced cognitive flexibility and cue-induced craving can result in a lapse or relapse by abstinent alcohol-dependent patients. Building on our findings, a mechanism-based intervention targeting the mGluR2 deficit in alcohol-dependent individuals is suggested. Using an established rat model of alcohol relapse *(45, 46)*, we showed that the mGluR2/3 agonists LY379268 and LY354740 dose-dependently reduced relapse-like drinking. Importantly, the repeated sub-chronic administration of an mGluR2/3 agonist did not lead to tolerance to its anti-relapse properties; a major concern reported with other approved anti-relapse compounds *(58, 59)*. However, mGluR2/3 agonists are unlikely candidates for the clinic because they do not discriminate well between the contributions of mGluR2 vs. mGluR3 receptors. In addition, these agonists are associated with adverse reactions *(54, 60)*. To overcome these limitations, PAMs selective for mGluR2 were recently developed. Here we show that repeated administration of the mGluR2 PAM LY487379 reduced relapse-like drinking in both male and female rats with no visible side-effects. Together with other data showing that PAMs can reduce alcohol-seeking behavior *(54)*, we conclude that PAMs for mGluR2 should be considered for clinical trials in alcohol-dependent patients.

Finally, we provide evidence for a candidate biomarker approach. [^18^F]-FDG PET, an index of regional glucose metabolism widely used in clinical diagnostics, can also be applied to biomarker development *(27)*. Several FDG-PET studies in alcohol-dependent patients have consistently reported decreases in overall brain glucose metabolism at rest *(61)*. This is also seen in alcohol-dependent rats that had long-term intermittent access to alcohol *(62)*. In the present study, alcohol-dependent rats also showed a decrease in whole brain glucose utilization, an effect that was also observed with the infralimbic cortex, in a more specific ROI analysis. Therefore, we conclude that FDG-PET studies in rat models of alcoholism may have translational utility, and can specifically be used for biomarker development. Following treatment with LY379268, no alterations in glucose levels were detected in alcohol-dependent rats with the infralimbic cortex, whereas in control rats reduced whole brain uptake was detected in this region. Non-responding to an mGluR2/3 agonist can be explained by the pronounced mGluR2 deficit in alcohol-dependent rats, especially in the corticostriatal system *(8)*. This FDG-PET finding most likely translates to humans, as we have also reported strong prefrontal mGluR2 expression deficits in deceased alcohol-dependent patients *(8)*. We conclude that alcohol-dependent individuals with an mGluR2 deficit do not respond in a detectable range to glucose update, but show a beneficial treatment effect in respect to craving and relapse. A likely interpretation of these findings is that the FDG response reveals a decreased sensitivity of mGlu2 activation due to decreased receptor expression, but that a sufficient receptor pool remains for this deficit to be rescued by agonist or PAM treatment.

Our multi-disciplinary series of experiments has at least two limitations. First, we show that a general loss of mGluR2 function did not affect ASST performance, a surprising finding given that we conclude here that mGluR2 is critically involved in cognitive flexibility. For a general loss of mGluR2 function model we used Indiana Alcohol preferring (P) rats, which carry the mGluR2 cys407* point mutation, preventing expression of functional mGluR2 receptors in the entire brain *(24, 63)*. Indiana P rats did not show any impairment in ASST performance compared to their respective control rat line, the Indiana non-preferring (NP) rats. This finding suggested that a general loss of mGluR2 function does not affect ASST performance. One possible explanation could be that mGluR3 can compensate for the loss of mGluR2 function in this rat line, as both receptors can act as presynaptic auto-inhibitors in neurons *(64, 65)*. Second, we used a neuron targeted viral mGluR2 knockdown in the infralimbic mPFC of CamKII-Cre rats to prevent general compensatory molecular mechanisms. However, we did not achieve an exclusive co-localization of Cre with CamKII neurons in the infralimbic cortex of CamKII-Cre rats. There was therefore not a complete neuronal restriction of the Cre-dependent AAV expression, which may explain why we observed only a partial rescue in the ASST.

Combined with other reports *(25, 26, 44, 54)*, our preclinical results provide support for mGluR2 as a molecular target for treating reduced cognitive flexibility, craving and relapse responses in alcohol-dependent patients. In terms of further clinical translation, we propose the following key steps. First, we suggest performing an experimental medicine trial in alcohol-dependent patients in order to demonstrate improved cognitive flexibility in response to a single administration of an mGluR2 PAM. Such a trial would benefit from an enrichment strategy based on the here described FDG-PET biomarker. Second, we suggest performing a cue-elicited craving study in alcohol-dependent patient in the MRI scanner in order to demonstrate normalized functional connectivity in brain areas known to be involved in neuronal cue reactivity following a single application of an mGluR2 PAM *(66–68)*. In the case that both proposed human experimental studies yield positive results, an RCT for testing the anti-relapse properties of a given mGluR2 PAM is indicated.

## Supporting information

Spplemental materials and methods

## Acknowledgments

We would like to thank Miriam Schneider and Anja Göpfrich for sharing their ASST protocol and help implementing it. Furthermore, we would like to thank Elisabeth Röbel, Sabrina Koch, Ana Gallego-Roman, Konstantin Wagner, Natalie Hirth, Batool Alharastani and the Thomas Kuner lab for technical assistance. Financial support for this work was provided by the Bundesministerium für Bildung und Forschung (BMBF) funded ERA-NET program: Psi-Alc (FKZ: 01EW1908), the BMBF-funded SysMedSUDs consortium (FKZ: 01ZX1909A), and the Deutsche Forschungsgemeinschaft (DFG, German Research Foundation) – Project-ID 402170461 – TRR 265 *(69)*, and the European Union’s Horizon 2020 program (668863-SyBil-AA)

## Author Contributions

M.W.M., S.P. W.H.S and R.S. designed research; M.W.M., S.P., C.R., V.V., J.B., R.H., M.L.M., E.R., A.C.H., G.K., O.B.H. and N.M., performed research; M.W.M., S.P., C.R., M.L.M., and W.H.S. analyzed data; R.L.B., H.E., B.N., K.S., and D.B. provided critical infrastructure or specific (transgenic)rat lines; M.W.M., S.P., R.S.,and W.H.S. wrote the paper.

