## Supplementary material for "A common molecular mechanism for cognitive deficits and craving in alcoholism": Spplemental materials and methods

### Supplementary Materials and Methods

#### Animals

Male Wistar rats (Charles River, Germany; initial weight 250 – 300g; n = 130), Indiana alcohol-preferring (RGD Cat# 634380, RRID:RGD_634380) and non-preferring rats (initial weight 250 – 300g, P: n = 12; NP: n = 14 *(63)*, bred at the animal production core facility at Indiana University, and CamKII-Cre transgenic rats (initial weight 250 - 300g; n = 28), bred on a Sprague-Dawley genetic background in the breeding facility of the CIMH Mannheim were housed in groups of four under a 12 h light/dark cycle with food and water available *ad libitum* in home cages. All behavioral testing was performed during the light phase (6:00 A.M. to 6:00 P.M.), 5 d per week and conducted in accordance with the EU guidelines for the care and use of laboratory animals and were approved by the local animal care committee (Regierungspraesidium Karlsruhe, Karlsruhe, Germany).

#### Materials

96% Alcohol from Sigma-Aldrich (24105-5L-R, Schnelldorf, Germany) was used for chronic intermittent alcohol exposure as well as for operant alcohol-seeking experiments. Analox Alcohol reagent kit (GMRD-113) from Campro Scientific (Berlin, Germany) was used to determine blood alcohol concentrations during alcohol vapor exposure. Casein pellets (Bio Serve Dustless Precision Pellets, Bilaney, Kent, UK) were used as reward during Attentional Set Shifting Task. Orange oil (Oleum Aurantii dulcis) was from Caelo. RNAscope® Fluorescent Multiplex kit (320850), RNAscope® Probe – Rn-Grm2 (317751) and RNAscope® Probe – EYFP-C2 (312131-C2) were from Advanced Cell Diagnostics. Primary antibodies: monoclonal mouse anti-GAD67 and mouse anti-NeuN (Millipore Cat# MAB5406, RRID:AB_2278725 and

Millipore Cat# MAB377, RRID:AB_2298772) and monoclonal mouse anti-CaMKII (Thermo Fisher Scientific Cat# MA1-048, RRID:AB_325403). Secondary antibody: AlexaFluor 555-labeled donkey anti-mouse (Thermo Fisher Scientific Cat# A-31570, RRID:AB_2536180). TOTO-3 Iodide (Thermo Fisher). Western Blot materials: cOmplete mini protease inhibitor (Roche Diagnostics, Mannheim, Germany); BioRad Protein Assay (Bio-Rad Laboratories, Munich, Germany); protein ladder (Chameleon® Duo Li-Cor, Lincoln, NE, USA); 1x Tris Glycine SDS Running Buffer (Invitrogen, Carlsbad, CA, USA); Novex™ Tris-Glycine Transfer buffer (Invitrogen); Odyssey® Blocking Buffer/ PBS (Li-Cor); primary antibodies for western blot: anti-metabotropic glutamate receptor 2 antibody (1:1000, Abcam Cat# ab15672, RRID:AB_302021, MA, USA) and rabbit anti-beta actin (1:3000, Cell Signaling Technology Cat# 4970, RRID:AB_2223172, Danvers, MA, USA). Secondary antibodies for western blot: IRDye 800 CW Donkey Anti-Mouse IgG (1:1000, LI-COR Biosciences Cat# 926-32212, RRID:AB_621847) and IRDye 680LT Donkey Anti-Rabbit IgG (1:1000, LI-COR Biosciences Cat# 926-68023, RRID:AB_10706167). For genomic DNA extraction the NucleoSpin® 8 / 96 Tissue kit (Macherey – Nagel, Düren, Germany) was used. The Grm2 cys407* SNP (c.1221C>A, p.Cys407*) was detected by a custom TaqMan® SNP Genotyping Assay (Assay ID: AHGJ96C, Applied Biosystems, Carlsbad, USA). For the GTP-γ-S Assay GDP (Sigma Aldrich), [35S]GTP-γ-S (PerkinElmer) and LY379268 (mGluR2 agonist, Tocris Bioscience) were used. [35S]GTP-γ-S signals were detected using FUJI imaging plates (Storage Phosphor Screen BAS-IP SR2025 Screen, GE Healthcare Life Sciences).

**Generation of CamKII-Cre transgenic rat**

CamKII-Cre transgenic rats were generated using a random integration BAC-approach (Fig 2A) A similar BAC was previously used to generate transgenic mice *(71)*, however we used a modified targeting vector for homologous recombination in Escherichia coli *(72)*, harboring the codon improved Cre recombinase only. The iCre cassette was inserted into the endogenous ATG of the CamKII-alpha gene. Linearized BAC DNA was purified and microinjected into the pronucleus of Sprague Dawley rat oocytes according to published protocols *(73)*. Transgenic founder rats were identified by PCR genotyping of tail tips and positive animals were bred on a Sprague-Dawley genetic background.

**Grm2 *407 genotyping**

A recent report from *(74)* indicates that the Grm2 cys407* SNP observed in Indiana P rats can also occur in wild type Wistar rat colonies. Therefore, in the present study Wistar rats and transgenic Sprague Dawley rats were screened for this premature stop-codon mutation and SNP carriers were excluded from all experiments.

Tissue for Grm2 cys407* genotyping was obtained from the animals by tail biopsy. Genomic DNA was isolated using the NucleoSpin® 8 / 96 Tissue kit (Macherey – Nagel, Düren, Germany) according to the manufacturer’s protocol. The Grm2 cys407* SNP (c.1221C>A, p.Cys407*) was detected by a custom TaqMan® SNP Genotyping Assay (Assay ID: AHGJ96C, Applied Biosystems, Carlsbad, USA) on an ABI QuantStudio 7 Flex RT-PCR system with QuantStudioTM Real-Time PCR software (20 µl reaction volume containing 10 ng genomic DNA, 55 cycles of 95 °C for 15 sec and 57 °C for 30 sec). Only homozygous wild type allele carriers were used for further experiments. We found that ~20% of transgenic Sprague Dawley and wildtype Wistar rats were heterozygous for the cys407* SNP, none were homozygous for cys407* and ~80% were homozygous wildtype allele carriers.

**Generation of a Cre-inducible mGluR2 knockdown virus**

In order to generate a Cre-inducible mGluR2 knockdown AAV, we cloned a construct adapted from the conditional by inversion (COIN) strategy *(75)*. A commercial siRNA sequence was used to construct the shRNA specifically targeting mGluR2 rat mRNA (siRNA ID: s127825, Silencer® Select Pre-Designed, Ambion, Thermo Fisher). To prevent Cre-independent unspecific shRNA expression, both the sense and antisense sequences were separated from each other as can be seen in Figure 2E. Only after Cre-recombination, the lox71 and lox66 flanked sequence flips irreversibly and a functional shRNA sequence is expressed under control of U6 promoter; eYFP is expressed under control of EF1α promoter, respectively DNA sequences of the cloned vectors can be obtained upon request.

**Luciferase reporter plasmid**

In order to verify the knockdown efficiency of the newly generated Cre-inducible shRNA AAV vector, a firefly luciferase reporter plasmid was cloned. For this purpose, the plasmid pMIR-REPORT™(ThermoFisher) was used and a target site for $mGluR_2$ was inserted.

**AAV production**

A standard protocol was used for AAV production *(76)*. Briefly HEK293 cells were transfected with three helper plasmids (pFdelta6, pRV1 and pH21) and the shRNA containing AAV plasmid by calcium phosphate precipitation. Sixty hours after transfection cells were harvested and purified by heparin columns.

#### Stereotaxic virus injections

For AAV (n = 28 CamKII-Cre rats) and lenti virus injections (n = 32 Wistar rats), rats were anesthetized with isoflurane and placed in a Kopf stereotaxic frame. Bilateral injections of 0.5µl of AAV or 1.2µl lenti virus into the infralimbic mPFC (AP: +2.9, ML: +/- 0.5, DV -5.1) according to *(38)* were performed using a WPI microinjection pump through a 33 gauge beveled needle at a rate of 200 nl/min. Behavioral testing was started four weeks (AAV) or two weeks (lenti virus) after virus injection.

#### Behavioral Assays

#### Operant alcohol self-administration

##### *Apparatus*

Operant alcohol-seeking experiments were performed in operant chambers (MED Associates) enclosed in ventilated sound-attenuating cubicles. The chambers were equipped with a response lever on each side panel of the chamber. Responses at the appropriate lever activated a syringe pump that delivered a ∼30 µl drop of fluid into a liquid receptacle next to it. A light stimulus (stimulus light) was placed above each response lever of the self-administration chamber, activated only upon correct lever responses. An IBM-compatible computer controlled the delivery of fluids, presentation of stimuli, and data recording.

#### *General operant alcohol self-administration procedure*

Alcohol self-administration training and testing sessions were performed 3 h after beginning of the dark phase, 5–6 d per week. Animals were trained to self-administer 10% (v/v) alcohol in daily 30 min sessions without prior sucrose-fading procedures. During the first 3 d of training, the animals were kept water-deprived for 18 h per day.

#### *Operant cue-conditioning*

The animals were trained to associate the availability of alcohol with the presence of specific discriminative stimuli, using a combination of discriminative (olfactory) and contingent (visual) cues *(8, 77, 78)*. Two stimuli were used to predict the availability of 10% (v/v) alcohol and were presented during each daily 30 min conditioning session. An orange odor served as an olfactory contextual stimulus for alcohol and was generated by the application of four to six drops of the orange extract onto the bedding material in the operant chamber before the start of each session. This stimulus was present throughout the whole session. In addition, a discrete visual stimulus was presented after correct responses resulting in alcohol delivery (left lever). As visual stimulus a 5 s blinking light was used, which was activated after a correct response at the active lever and was therefore directly connected to alcohol availability. The 5 s period served as a “time out”, during which responses were recorded, but did not lead to reinforcer delivery. For the stimulus-conditioning training, the animals had to complete 15 sessions with the two conditioned stimuli (CS). During the stimulus conditioning training responses at the right lever were recorded, but did not result in reward delivery (inactive lever). To determine the baseline of self-administration responses, the mean over the last three self-administration sessions was calculated.

#### *Extinction training*

After successful completion of the stimulus conditioning phase, all animals underwent daily 30 min extinction sessions for 5 consecutive days, which was sufficient to reach an extinction criterion of < 10 % of baseline activity at the active lever per session. During extinction sessions, both levers were extended without presentation of the olfactory CS (orange odor). Responses at the previously active lever activated the respective syringe pump; which did not result in reinforcer delivery or presentation of the discrete CS (blinking light).

#### *Cue-induced reinstatement test*

The animals were presented with the same conditioned stimuli (CS) as during the conditioning phase. Responses at the left (active) lever resulted in the presentation of the visual blinking light stimulus and activation of the syringe pump; which did not result in reinforcer delivery.

### Attentional Set Shifting Task (ASST)

#### *Test apparatus*

The test apparatus was made of dark grey PVC consisting of a small compartment (20 cm x 40cm x 40 cm) adjacent to the test compartment (40 cm x 50 cm x 40 cm). The two compartments were separated by a sliding door (width 20 cm). Two small ceramic bowls (diameter 7 cm, depth 4 cm) were positioned into the test compartment 16 cm apart from each other, separated by a divider. The two bowls were filled with different digging materials and/or were differently scented. A casein pellet (Bio Serve Dustless Precision Pellets, Bilaney, Kent, UK) served as a reinforcer and was deeply buried in one of the bowls. Rats were trained to dig in the bowl to retrieve the reinforcement. The presence or absence of the reinforcement pellet in the digging bowl was signaled by either an olfactory (odor) or a visual-tactile cue (shape and tactile quality of digging medium).

#### *Habituation*

Animals were familiarized with the food reinforcer, the ceramic bowl, and the different digging materials in their home cage prior to testing. During 1-2 nights prior to the test, the pots were filled with home cage bedding, and several casein pellets were presented at the top of the bedding as well as deeply buried within the bedding. The pots were rebaited regularly and left in the home cage overnight. The following night, the digging media used for the pretraining period, simple and compound discrimination task were baited and similarly placed in the home cage. On the second day of habituation, two familiar animals were placed into the test apparatus and were allowed to explore it freely for 15 min. On the third day of habituation, each rat was placed into the test apparatus individually for another 15-min habituation period. During the entire habituation and testing period, the animals were maintained at approximately 90% of their free-feeding weight (12 g/rat/day). Food restriction started one week prior to testing.

#### *Testing procedure*

The testing procedure was adapted from *(18, 19).* After habituation, all animals underwent a pretraining schedule, during which the animals had to retrieve the reward from empty pots and subsequently from pots filled with digging medium. First, the reward was placed on top of the digging medium and was subsequently buried deeper into the medium in further trials. Each trial within the pretraining schedule was repeated until the reward was retrieved. The rats had to retrieve the pellet five times within two minutes, followed by four pellet retrievals within one minute. As soon as the rat retrieved the reward pellet or the trial time expired, the animal was gently pushed back into the starting area. The pots were rebaited during the inter-trial-interval (ITI, 30s). During this time, the rat had to wait inside the starting area until the sliding door was lifted for the next trial. The digging medium from the pretraining procedure was not used during further testing.

Eight common spices and media were used for all discrimination tasks, which are listed in Table 1. The digging media were intermixed with powdered casein pellets to avoid olfactory detection of the pellet in the bowl. During all testing sessions, a criterion of six consecutive correct trials was used for successful learning (trials to criterion).

**Table 1: Examples of odor-medium pairs used for ASST**

| Digging medium | Digging medium | Odor 1 | Odor 2 |
|---|---|---|---|
| Seramis | | | |
| Colored silica sand | Hamster bedding | Cumin | Capsicum |
| Beech chipping | Rough stones | Nutmeg | Basil |
| Straw pellets | Pine bark | Thyme | Dill |
| Cork granules | Black silica sand | Rosemary | Curcuma |

During the ASST the animals were tested in the following subtasks:

Simple Discrimination (SD):

Two bowls containing different media but scented with the same odor were presented to each rat. The visual/tactile stimulus indicated the position of the reward (Medium 1 (M1)).

Compound Discrimination (CD):

For the compound discrimination, an additional odor was introduced and used together with the two previously used media and the previously used odor. The previously baited digging medium used during SD (M1) also indicated the location of the reward during CD, independent from the presented odors.

Compound Discrimination reversal (CDrev):

The previously learned rule was reversed. The previously baited medium was not baited, but the second medium was baited instead (M2).

Compound Discrimination repetition (CDrep): A repetition test of CDrev.

Intradimensional shift (IDS):

Introduction of a new set of complex stimuli. The animals had to discriminate the baited from the unbaited bowl by using the same perceptual dimension (digging medium) as in the previous testing.

Extradimensional shift (EDS):

A new set of stimuli was introduced. However, now the previously irrelevant dimension predicted the reward. Therefore not the type of digging material predicted the reward, but the odor was relevant to obtain the pellet.

If an animal stopped responding for several trials during a test session, it was returned to the homecage for up to 1 h before resuming the test again. In this case, the sum of the number of trials was taken.

**Delay discounting test**

Rats were trained in operant conditioning chambers (MED Associates) to self-administer a 0.2% saccharin solution (Sigma-Aldrich). The protocol was adapted from *(20)*. Animals were trained 5 days a week, one session per day containing 50 free choice trials. Within every trial there was a response window of 20s followed by an intertrial interval of 65s; no response was scored as trial omission. The intertrial interval was automatically adjusted to make sure that a new trial always starts exactly 65s after a response had been made.

After habitation to the operant boxes, the initial training stage began where two levers were presented with identical reward size (30 µl/trial of 0.2% saccharin solution) until rats had made >100 lever presses/30min. In the next stage, reward size discrimination training was conducted. Here, one lever was randomly assigned as high reward lever (90 µl/trial) and the second as low reward lever (30µL/trial). The behavior of the animals was observed during each session until the mean preference for the high reward lever was >85% of total trials. In the following discounting task, a time delay was introduced to the high reward lever (i.e. before the higher reward was presented). The timepoints for the delays were (0, 6, 12, 24, 36, 48, and 60 seconds); each delay was introduced on a separate day sequentially. In the delay discounting task, 10 forced choice (FC) trials were introduced at the beginning of each session, 5 on each lever in a random order, to make the rats aware of their upcoming choices and preventing passive behavior (i.e., inactivity). Behavioral experiment results were recorded via Med PC software. The software was programmed to record the number of trials that the rat chose in each session, whether the choice was for the high reward, low reward or neither (i.e. omission).

**Alcohol vapor exposure**

For the chronic intermittent ethanol (CIE) vapor/ air exposure we used a protocol adapted from *(21)*. Briefly, pumps (Knauer) delivered alcohol into electrically heated stainless-steel coils (60°C) connected to an airflow of 18 L/min into glass and steel chambers (1 × 1 × 1 m). For the next 8 weeks rats were exposed to five cycles of 14 h of alcohol vapor per week (0:00 A.M. to 2:00 P.M.) separated by daily 10 h periods of withdrawal. Twice per week, blood (~20 μl) was sampled from the lateral tail vein, and blood alcohol concentrations were determined using an AM1 Analox system (Analox Instruments). After the last exposure cycle the rats stayed withdrawn for two weeks before further behavioral testing.

**Long-term voluntary alcohol consumption with repeated deprivation phases**

After two weeks of habituation to the animal room, rats were given ad libitum access to tap water and to 5%, 10% and 20% alcohol solutions (v/v) concurrently. Spillage and evaporation were minimized by the use of special bottle caps. With these bottle caps, the alcohol concentration remains constant for at least one week *(79)*. The position of the bottles was changed weekly. The first deprivation period was introduced after eight weeks of continuous alcohol availability. After a deprivation period (between 2-3 weeks), rats were given access to alcohol again and five more deprivation periods were introduced in a random manner, i.e., the duration of drinking and deprivation phases was irregular, i.e. approximately 4 ± 1 week and 2 ± 1 week, respectively in order to prevent behavioral adaptation *(58, 80)*. The long-term voluntary alcohol drinking procedure, including all deprivation phases, lasted a total of 12 months.

In order to study the effects of different drugs in comparison to vehicle, rats were divided into groups (n = 8/10) in such a way that the mean baseline total alcohol intake was approximately the same for each group. Baseline drinking was measured daily for one week. After the last day of baseline measurement, the alcohol bottles were removed from the cages, leaving the animals with free access to food and water for the complete deprivation period.

In the current study, we tested three drugs. The chosen drug administration scheme was a repeated dosing scheme (i.e., five doses across three days, starting Monday night, followed by administration twice daily on Tuesday and Wednesday) based on an established paradigm used in our previous ADE studies to test alcohol relapse behavior *(58, 81)*.

### Neuroanatomical Experiments

#### *Injection site mapping*

In order to validate virus injection sites, all virus injected animals were deeply anesthetized with isoflurane and intracardially perfused with 100 ml of 1× phosphate buffered saline (PBS) followed by 200 ml of 4% paraformaldehyde (PFA) in 1x PBS. Brains were collected and postfixed for 24 h at 4°C in 4% PFA in 1x PBS. Next 55 µm coronal sections were cut using a vibrating blade microtome (Leica Microsystems), mounted on microscope slides using Immu-Mount (Fischer Scientific) and validated by recording GFP fluorescence at the injection sites using a Zeiss Axioskop 2 plus microscope with a 2.5× lens.

#### *Immunohistochemistry procedures*

In order to characterize the cells expressing Cre-inducible mGluR2 knockdown AAV, 3 CamKII-Cre rats received bilateral injections of the knockdown AAV into the infralimbic mPFC. Four weeks after virus injection rats were perfused and postfixed as described above. Sixty µm coronal sections were cut using a vibrating blade microtome (Leica Microsystems) and collected in 1x PBS. Immuno-labeling for NeuN, GAD67 and CamKII were performed as previously described *(78)*. Briefly, sections were washed in 1x TBS and incubated in blocking solution (7.5% donkey serum, 2.5% BSA in 1× TBS with 0.2% Triton X-100) for 1h at RT. Sections were either incubated in anti-NeuN primary antibody (mouse, 1:500, MAB377 Millipore) or anti-GAD67 (mouse, 1:1000, 1G10.2 MAB540, Millipore) or anti-CamKII (mouse, 1:500, 6G9 MA1-048, Pierce Biotechnology) in blocking solution for 24 h at 4°C. After washing with 1x TBS sections were incubated in secondary antibody solution containing AlexaFluor 555-labeled donkey anti-mouse (1:200, Invitrogen) in 1x TBS containing 0.2% Triton X-100 for 2 h at RT. Sections were washed with 1x TBS, counterstained with TOTO-3 Iodide (1:2000 in PBS, Thermo Fisher) and mounted as described above.

***RNAscope fluorescent in-situ hybridization***

To quantify the knockdown efficiency of the Cre-inducible mGluR2 knockdown AAV on mRNA level, three male CamKII-Cre rats were injected with the control AAV expressing shUnc and three rats were injected with the mGluR2 knockdown AAV. Four weeks after AAV injection the animals were rapidly decapitated. Brains were removed, frozen in isopentane (-50°C), and kept at -80°C until further processing. Brain slices of 20 µm thickness were cut on a cryostat and thaw-mounted onto Super Frost Plus slides (Thermo Fisher, Darmstadt, Germany). FISH analysis was performed using the RNAscope Multiplex Fluorescent Reagent Kit (Advanced Cell Diagnostics, Newark, USA; Probes Rn-Grm2 and EYFP-C2) according to the manufacturer's instructions (freshly frozen tissue).

Brain sections containing the mPFC were examined by confocal microscopy using a Leica TCS SP5 microscope (Leica Microsystems, Wetzlar, Germany) equipped with a 63x HCX PL APO (1.45 NA) objective. Three images were acquired at random positions in the infralimbic cortex of each hemisphere. Two brain slices were examined per animal.

Co-localization of mGluR2 and EYFP signals were analyzed using the cell counter macro in ImageJ. The specificity of the FISH signal was verified by co-localization with DAPI. Next, the mean grey value of mGluR2 signals were determined by manually drawing ROIs containing the perinuclear mGluR2 signals of each cell. For control as well as knockdown AAV injected animals the mGluR2 mean grey value ratio was calculated: $\frac{mGluR2^{+}\ EYFP^{+}}{mGluR2^{+}\ EYFP^{-}} = mGluR2\ ratio.$

***Spine density analysis – Golgi impregnation***

For Golgi impregnation, the animals were perfused, and brains were postfixed as described above. The right hemisphere of the brains was used for Golgi impregnation. Golgi-impregnation was performed according to the Golgi-Cox procedure using the Rapid GolgiStain reagent (FD NeuroTechnologies, USA) as described previously *(82)*. 120 µm coronal serial sections were cut using a vibratome (VT 1000E, Leica, Germany). The mPFC and NAc areas were identified using Paxinos and Watson (1998). Analysis of dendritic spines was conducted in a blinded procedure. Only secondary and tertiary dendrites were evaluated, which displayed no breaks in their staining and were not obscured by other neurons or artifacts. Only one segment per individual dendritic branch and neuron was chosen for the analysis, as recently described *(83)*. Quantitative three-dimensional analyses of dendritic fragments with their spines were conducted using NeuroLucida (MBF Bioscience, USA) controlling the x–y–z-axis of the microscope (Axioscop Imaging, Zeiss, Germany) and a microscope-mounted camera (AxioCam, Zeiss, Japan). The three-dimensional reconstruction was done using a 100× objective (NA: 1.4; oil immersion) and the NeuroLucida system. Spine densities were calculated from the reconstructed dendrites with the help of NeuroExplorer (MBF Bioscience, USA), as described previously in detail *(84)*. At least 20 dendrites (n = 20) per region and animal were reconstructed. Per group, five individual brains were analyzed (N = 5). The N values for the statistical analysis were based on animal numbers (N) and not on numbers of analyzed elements (n). For statistical evaluation (unpaired t-test), the statistical software package GraphPad Prism 5 (Graphpad Software, San Diego, CA, USA) was used.

**Biochemical Assays**

***Dual-Luciferase assay***

In order to quantify the knockdown efficiency of the Cre-inducible mGluR2 knockdown AAV, a dual luciferase assay was performed. To induce the Cre-mediated switch of the floxed mGluR2 knockdown cassette, the plasmid was transformed into EL350 E.coli cells *(85)*. For the quantification of mGluR2 knockdown efficiency, the mGluR2 target sequence was inserted into a pMIR-REPORT miRNA Expression Reporter Vector System (Thermo Fisher, Waltham, MA, USA). Both the control AAV construct (shUnc) and the recombined mGluR2 knockdown AAV construct were each co-transfected with the pMIR-REPORT vector (containing the mGluR2 knockdown target site and Firefly luciferase) and pSV40-Renilla containing Renilla luciferase for normalization into HeLa cells (jetPRIME™, Polyplus transfection, Illkirch, France). Three technical replicates were performed for each plasmid. Transfected HeLa cells were incubated for 48h at 37°C and 5% CO2 in Dulbecco´s Modified Eagle Medium (DMEM) + GlutaMax-I (Invitrogen) supplemented with 10% fetal calf serum (FCS) (Invitrogen) and 1% Streptomycin/Penicillin (Invitrogen). After 48h, cells were washed with 1x PBS and placed on ice. PBS was replaced with 1× Passive lysis buffer (Promega), and cells were harvested using cell scrapers (VWR). The cell suspension was transferred into a pre-cooled Eppendorf tube and centrifuged for 5 min at 4°C at 13200 rpm. Ten µl of each lysate was then analyzed using VICTOR 1420 Multilabel Counter (PerkinElmer, Hamburg, Germany) with automatic LAR II (Firefly luciferase) and Stop & Glo Reagent (Renilla luciferase) injection. All Firefly luciferase signals were normalized to Renilla luciferase signals.

***Western blot***

Protein levels of mGluR2 were examined within the NAc Shell of control or Cre-inducible knockdown AAV injected male CamKII-Cre rats (n = 8/group).

NAc shell brain tissue was micropunched as previously described *(8)*. Brain tissue was transferred to a lysis buffer: 100 mM Tris HCl pH 8 and 20 mM EDTA containing cOmplete mini protease inhibitor (Roche Diagnostics, Mannheim, Germany). Ultrasonic lysis was performed using an ultrasonic device (Branson Sonifier 250, Danbury, CT, USA). Protein concentrations were analyzed using BioRad Protein Assay (Bio-Rad Laboratories, Munich, Germany) and visualized using a microplate spectrophotometer (PowerWave XS, BioTek Instruments, Bad Friedrichshall, Germany).

Next samples were mixed with Laemmli 2× buffer (4% sodium dodecyl sulfate, 10% β-mercaptoalcohol, 0.004% bromphenol blue and 0.125 Tris-HCl pH 6.8). An amount of ~10 µg of total protein as well as 5 µl of pre-stained protein ladder (Chameleon® Duo Li-Cor, Lincoln, NE, USA) was loaded and separated on Novex™ WedgeWell™ 4-12% Tris-Glycine Gels using 1× Tris Glycine SDS Running Buffer (Invitrogen, Carlsbad, CA, USA). Proteins were then transferred to nitrocellulose membranes (Protran BA85, Cat no. 10401196, GE Healthcare Life

Sciences Whatman$^{TM}$) in a blotting chamber (X Cell II$^{TM}$ Blot Module, Invitrogen) using Novex$^{TM}$ Tris-Glycine Transfer buffer (invitrogen). Membranes were then incubated in Odyssey® Blocking Buffer in PBS (Li-Cor) for 1h at RT and probed with mouse anti-mGluR2 antibody (1:1000, mG2Na-s, ab15672, Abcam, MA, USA) and rabbit anti-beta actin (1:3000, #4970, Cell Signaling Technology, Danvers, MA, USA) diluted in Odyssey® Blocking Buffer in PBS for 2 days at 4°C. Blots were then washed with 1× PBS followed by 1× PBS-T (containing 0.5% Tween20). Next, blots were incubated for 2h at RT in secondary antibody solution containing IRDye 800 CW Donkey Anti-Mouse IgG (1:1000, LI-COR®, 926-32212) and IRDye 680LT Donkey Anti-Rabbit IgG (1:1000, LI-COR®, 926-68023) in Odyssey® Blocking Buffer/PBS, followed by washing steps with 1× PBS, 1× PBS-T and aqua bidest. Signals were detected using an ODYSSEY® CLx234 (LI-COR®, Lincoln, NE, USA) fluorescent imaging system. The signal intensity was quantified using Image Studio$^{TM}$ 235 software (LI-COR®, Lincoln, NE, USA) by calculating the ratio of mGluR2 signal normalized to ß-actin.

***[35S] GTP-γ-S autoradiography***

In order to detect LY379268-stimulated G-protein coupling after mGluR2 receptor knockdown, male Wistar rats received either an injection of the mGluR2 shRNA knockdown virus or control AAV (shUnc) into the infralimbic mPFC (n = 4/group). Four weeks after AAV injection, the animals were killed by rapid decapitation, brains were removed, frozen in isopentane (−50°C), and kept at −80°C until further processing. Twelve µm coronal brain sections were cut throughout the NAc using a cryostat (Bregma +1.7 mm to +0.70 mm). Sections were rinsed for 10 min at RT in 50 mM Tris–HCl pH 7.42, 3 mM MgCl2, 0.2 mM EGTA, 100 mM NaCl followed by incubation for 20 min at RT in 50 mM Tris–HCl pH 7.42, 3 mM MgCl2, 0.2 mM

EGTA, 100 mM NaCl, 0.1 % BSA containing 2 mM GDP (Sigma-Aldrich). Sections were then incubated for 2 h at RT in the same buffer containing 2 mM GDP, 80pM [35S]GTP-γ-S (PerkinElmer), and 10 μM LY379268 (mGluR2 agonist, Tocris Bioscience). Incubation was stopped by washing the slides for 3×2 min in ice-cold buffer (50 mM Tris–HCl pH 7.4) followed by a dip in ice-cold deionized water.

Sections were dried under a stream of cold air and exposed against FUJI imaging plates (Storage Phosphor Screen BAS-IP SR2025 Screen, GE Healthcare Life Sciences) for several hours and scanned using a phosphorimager (Typhoon FLA 700, GE Healthcare, Germany). Densitometry analysis was performed using the MCID program (MCID Image Analysis Software Solutions for Life Sciences); measurements were compared against standard curves generated using [14C]-Microscales (Amersham, GE Healthcare Life Sciences). Agonist-stimulated and baseline GTP-γ-S bindings were measured on adjacent sections.

### Electrophysiology

#### *Brain Slice Preparation*

Rats were anesthetized with isoflurane and then perfused with a 4°C sucrose-based cutting buffer containing (in nM) 65 sucrose, 87 NaCl, 25 $NaHCO_3$, 2.5 KCl, 1.25 $NaH_2PO_4$, 10 Glucose, 0.5 $CaCl_2$, 7 $MgSO_4$, and 0.01 Pyruvate. Coronal slices (300 μm) containing both mPFC and NAc were obtained using a vibrating microtome (Leica VT 1200S Vibratome). Brain slices were transferred to regular artificial cerebrospinal fluid (aCSF) containing (in mM) 125 NaCl, 25 $NaHCO_3$, 2.5 KCl, 1.25 $NaH_2PO_4$, 2 $CaCl_2$, 1 $MgCl_2$ and 10 glucose with an osmolarity of 315 to 320 mOsm. Brain slices were allowed to incubate at 34°C for 30 min and then were placed at RT before recordings started. All solutions were continuously oxygenated (95% $O_2$, 5% $CO_2$).

***Brain Slice Recordings***

Recordings were performed at 31°-33°C in a submerged chamber perfused at 1.5-2.5 ml/min with oxygenated standard aCSF (see above). For electrical stimulation borosilicate glass micropipette (< 3 µm tip diameter; 0.5 mm wall thickness, 2 mm outside diameter) prepared with a Flaming/ Brown micropipette puller P-97 (Sutter Instruments, Novato, CA), filled with 0.9% NaCl solution and was placed in the infralimbic mPFC, ~300 µm away from the NAc-shell. Borosilicate glass recordings pipette (0.15 mm walled, 1.5 mm outside diameter) prepared with a Flaming/ Brown micropipette puller filled with aCSF and were placed in NAc-shell. Field potentials evoked at a rate of 0.033 Hz were acquired with an EPC10 amplifier (HEKA, Lambrecht, Germany), interfaced to HEKA Patchmaster software. Data were sampled at 20 kHz and low filtered at 3 kHz. Stimulation intensity was set to evoke 80% of the maximum field potential amplitude. Excitatory postsynaptic potential (EPSP) amplitude was measured as the difference of 5 voltage values before the stimulation and the peak of the EPSP. EPSP amplitudes were normalized to the average of the 10 min baseline period. mGluR2 agonist LY379268 (100 nM) was bath-applied for 5 min via perfusion. Following the LY379268 application, perfusion was switched back to aCSF to washout the agonist.

**Magnetic resonance imaging (MRI) and positron emission tomography (PET)**

After completion of 7 weeks of CIE or air exposure (n=32), animals were scanned in a 9.4 T horizontal bore animal scanner (Bruker, Rheinstetten, Germany). The animals were anesthetized during the scan by 2 % isoflurane in a gas mixture (O2:air (3:7)). Respiration rate and body temperature were monitored during each scan session. A T2-weighted sequence, rapid acquisition with relaxation enhancement (RARE) was used: RARE factor = 8, repetition time/effective echo time = 6500/32.5 ms, averages = 2, matrix size = 256×256, FOV = 3.2×3.2 cm2, 58 slices, slice thickness = 0.5 mm, interslice spacing = 0.5 mm.

After the MRI scans, during the two weeks of abstinence, animals were transported in specialized transport boxes containing air filters to a different lab for further pharmacological PET studies.

Animals were allowed to habituate to the animal facility in Cologne for one week prior to PET scanning. Animals received an i.p. injection of 2 mg/kg (injection volume 1mL) of the mGluR2 agonist LY379268 30 min prior to i.p. injection of 500-700 µl [$^{18}$F]-FDG solution (~2 mCi). The time interval of LY379268 injection was aligned to procedures used in other labs. LY379268 application 30 min prior to behavioral performance has been shown to lead to reduced reinstatement in an animal model of alcohol dependence *(44)*. Furthermore, the effects of the agonist last at least 180 min *(8)*, which is sufficient for the timeline of the PET experiment. After the injection of [$^{18}$F]-FDG, rats were placed in their home cage for 45 min. For the subsequent PET scan, rats were anesthetized (5% isoflurane for induction and 2.0–2.5% for maintenance in O2/N2O (3:7)) and placed and fixed on an animal holder (medres®, Cologne, Germany) with a respiratory mask. PET scans were performed using a Focus 220 micro PET scanner (CTI-

Siemens®) with a resolution at the center of the field view of 1.4 mm. A 10 min transmission scan using a $^{57}Co$ point source for attenuation correction was followed by a 30 min emission scan. Each emission scan started precisely one hour after [$^{18}F$]-FDG injection. To monitor the animal's condition, breathing rates were measured and kept around 60/min by adjusting isoflurane concentration individually. In addition, the body temperature was monitored and kept at 37°C by a feedback-controlled system.

#### *[$^{18}F$]-FDG -PET data analysis*

Following Fourier rebinning, summed images (60-90 min post [$^{18}F$]-FDG injection) were reconstructed using the iterative OSEM3D/MAP procedure *(86)*, resulting in voxel sizes of 0.38 × 0.38 × 0.82 mm. Imaging data were analyzed using the imaging software tool VINCI 5.0 (Max Planck Institute for Metabolism Research; available at vinci.sf.mpg.de). MR images were matched to a standardized rat brain atlas *(87)*, served as anatomical templates, and facilitated the coregistration of PET images. The assignment and designation of brain areas were based on the brain atlas of Paxinos and Watson. Subsequently, PET image intensities were normalized by the use of the ratio normalization technique *(88)*. That is, the intensity was divided by the mean in the reference area (standard uptake value ratio (SUVR)). For that purpose, the lateral ventricles were chosen as reference area (SUVRv), as the cerebral glucose utilization was supposed to be reduced in alcohol-dependent animals compared to controls *(56)*, and affected by the LY379268 treatment in various brain regions. For each of the 16 animals (8 control, 8 alcohol-dependent animals), two [$^{18}F$]-FDG-PET images were available (resting state condition with and without prior LY379268 injection).

**Statistics**

All data are expressed as mean ± SEM. Alpha level for significant effects was set to 0.05. Operant alcohol-seeking between-group data and ADE experiments were analyzed using repeated measures ANOVA, followed by Newman-Keuls post hoc test (Statistica 10, Statsoft; RRID:SCR_015627), where appropriate. Cue-induced reinstatement within-group data were analyzed using two-tailed t-tests.

Data of ASST subtasks were analyzed using one-way ANOVA. Overall group differences across all ASST subtasks were analyzed using repeated measures ANOVA.

Spine density data were analyzed using two-tailed t-tests.

Luciferase Assay and RNAscope in-situ hybridization data were analyzed using two-tailed t-tests. Western blot data were analyzed using one-tailed t-tests because of the expected reduction in protein levels in the mGluR2 knockdown group. For GTP-γ-S binding assays, values were calculated as percent of baseline value in the same region and animal and expressed as percent stimulation (± SEM) and were statistically analyzed by one-way ANOVA.

To visualize activation patterns in the FDG-PET experiment of control and alcohol-dependent animals during resting state as well as regions of activations and deactivation after LY379268 injection in comparison to resting state glucose utilization, an unpaired and two paired t-tests were performed (t-maps, Figure 5 C columns 1-3). To correct for multiple testing with 19,536 brain voxels, a threshold-free cluster enhancement (TFCE) procedure with subsequent permutation testing, thresholded at $p < 0.05$, was used *(69, 89)* for all t-maps. Because TFCE values are arbitrary, color bars of TFCE maps were labeled with the original t-values, marked tTFCE, respectively.

**Supplementary Figures:**

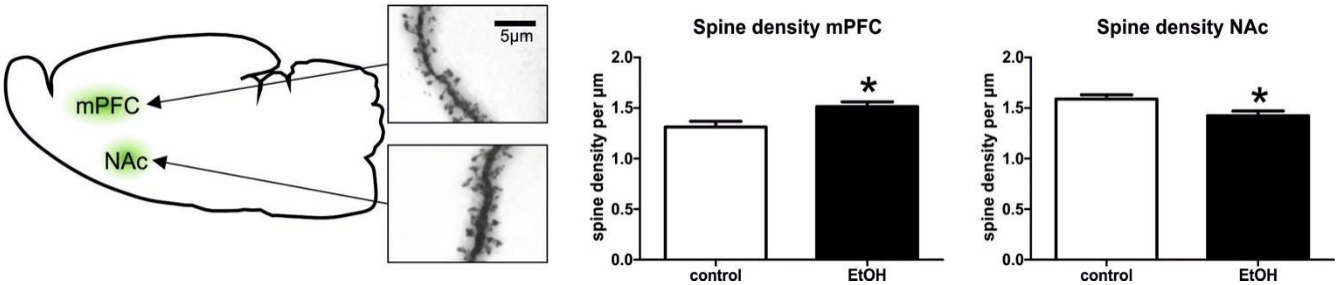


**Supplementary Figure 1: Spine density analysis in alcohol-dependent and control rats**. *Left:* Schematic representation and representative images of spine density analysis in mPFC and NAc. Spine density analysis revealed significant differences between alcohol-dependent and control rats (n = 5/group) in the mPFC (*middle*) and NAc (*right*).

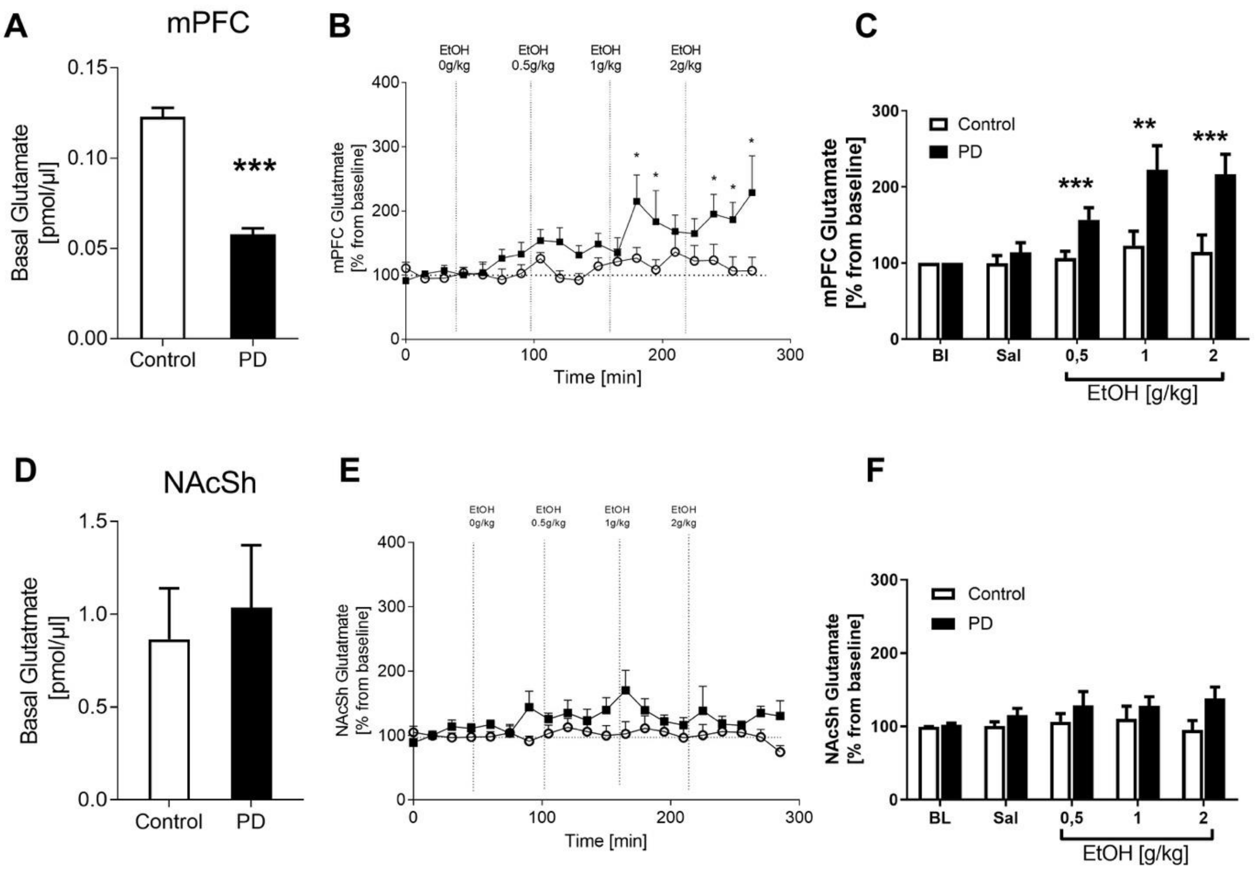


**Supplementary Figure 2: Glutamate microdialysis in the mPFC and NAcSh in freely moving alcohol-dependent and control rats. A)** The PD-mPFC has strongly reduced basal glutamate levels indicative of a hypo-glutamatergic status. **B&C)** Systemic administration of increasing doses of alcohol (0, 0.5, 1.0, and 2.0g/kg i.p) led to a robust dose-dependent increase in extracellular glutamate levels in alcohol-dependent rats with no changes in non-dependent rats. **D)** In the NAc shell we found a tendency towards increased glutamate baseline levels and systemic administration of increasing doses of alcohol (**E & F**) had no effect on extracellular glutamate levels neither in alcohol-dependent nor in control rats. * $p < 0.05$, ** $p < 0.01$, *** $p < 0.001$.

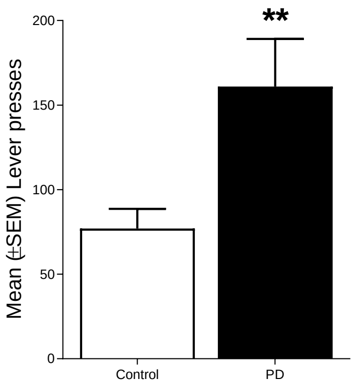


**Supplementary Figure 3: Alcohol self-administration of control and alcohol-dependent rats.** After seven weeks of CIE exposure, alcohol-dependent rats significantly increased their alcohol self-administration compared to controls. ** $p < 0.01$.

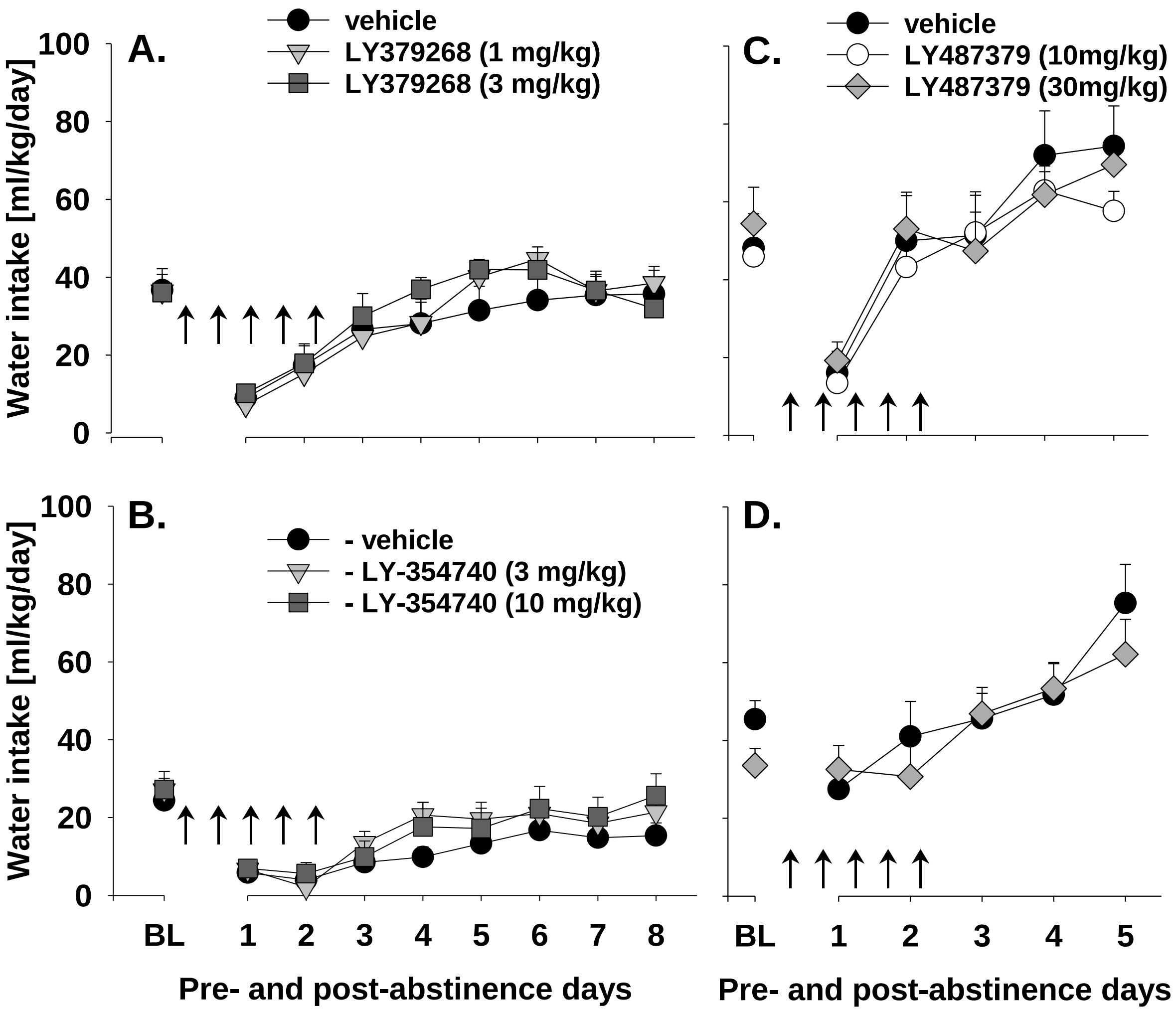


**Supplementary Figure 4.** Total water intake (ml/kg/day) before and after an alcohol deprivation period of three weeks. Arrows indicate the administration of (A) vehicle (n = 8), 1 mg/kg of LY-379268 (n = 8) and 3 mg/kg of LY-379268 (n = 8), (B) vehicle (n = 10), 3 mg/kg of LY-354740 (n = 8) and 10 mg/kg of LY-354740 (n = 7) in male rats and administration of either vehicle or LY487379 in both (C) male and (D) female rats [vehicle (n = 8 males, n = 10 females), 10 mg/kg of LY487379 (n = 7 males) and 30 mg/kg of LY487379 (n = 7 males, n = 8 females)]. Each animal was given total of 5 IP injections every 12 h. The last three days of alcohol measurements are given as baseline drinking – “BL”. All animals were re-exposed to alcohol immediately after the second drug administration. Data are presented as means ± S.E.M.

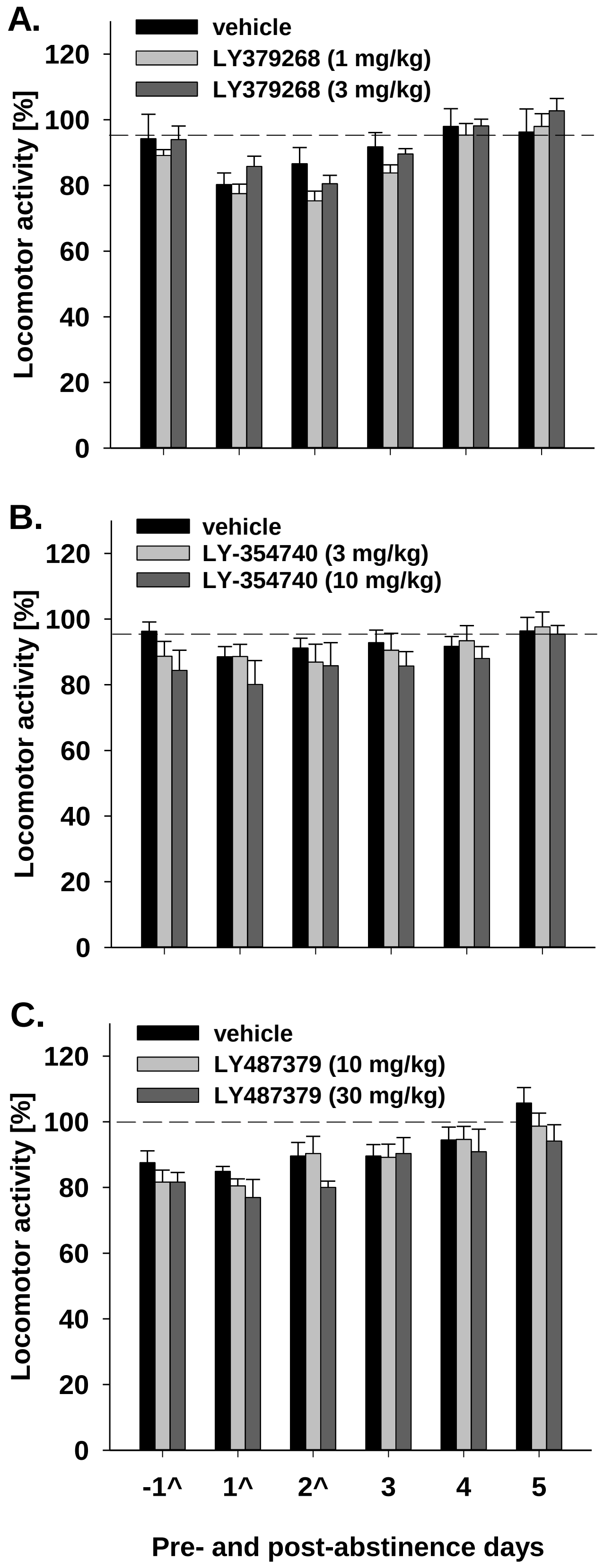
A.
vehicle
LY379268 (1 mg/kg)
LY379268 (3 mg/kg)
Locomotor activity [%]
0
20
40
60
80
100
120
B.
vehicle
LY-354740 (3 mg/kg)
LY-354740 (10 mg/kg)
Locomotor activity [%]
0
20
40
60
80
100
120
C.
vehicle
LY487379 (10 mg/kg)
LY487379 (30 mg/kg)
Locomotor activity [%]
0
20
40
60
80
100
120
-1^
1^
2^
3
4
5
Pre- and post-abstinence days

**Supplementary Figure 5.** Locomotor activity in (A) vehicle (n = 8), 1 mg/kg of LY-379268 (n = 8) and 3 mg/kg of LY-379268 (n = 8), (B) vehicle (n = 10), 3 mg/kg of LY-354740 (n = 8) and 10 mg/kg of LY-354740 (n = 7) and (C) vehicle (n = 8), 10 mg/kg of LY487379 (n = 7) and 30 mg/kg of LY487379 (n = 7) treated male rats. Locomotor activity is shown as 12-hour post-injection intervals of the animals' active phase. The percentage of each rat's locomotor activity during and after treatment days was calculated with respect to basal activity prior to treatment (dashed line). Each animal was given total of 5 IP injections every 12 h. Injection days are marked as "^". All animals were re-exposed to alcohol immediately after the second drug administration. Data are presented as means ± S.E.M.

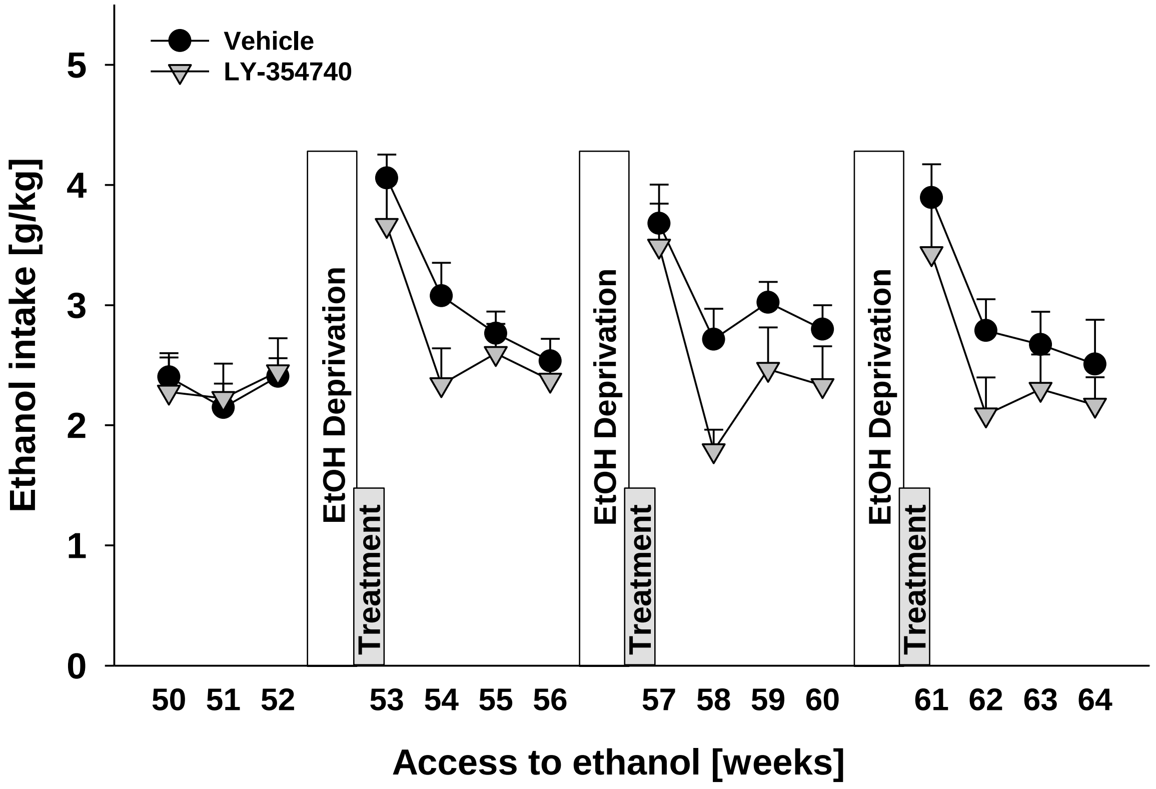


**Supplementary Figure 6.** Alcohol intake in either vehicle (n = 8) or 5 mg/kg of LY-354740 (n = 7) treated animal groups. Alcohol consumption was calculated in g of pure alcohol per kg of body weight per day and presented as the average weekly intake. Drug treatment started on the last day of the 7th alcohol deprivation, which was introduced following 52 weeks of intermittent access to alcohol. Each animal was given total of 5 IP injections every 12 h before and during three subsequent post-abstinence drinking phases (weeks 53, 57 and 61). Data are presented as means ± S.E.M.

**Supplementary Table 1 Statistics for operant alcohol-seeking behavior.** Results for repeated measures ANOVA and Newman-Keuls post hoc test are shown. Comparison between active and inactive lever was done using two-tailed paired t-test. Abbreviations: DF = degrees of freedom, F = F-value, p = p-value,  KD = knockdown, BL = self-administration baseline, EXT = extinction, RE = cue-induced reinstatement

| **Lever** | **Repeated measures ANOVA** | | | | | **Newman-Keuls post hoc test** | | | | | **t-test (comparison active/ inactive)** | |
|---|---|---|---|---|---|---|---|---|---|---|---|---|
| | **Test** | **DF** | **Effect** | **F** | **p** | **Within group comparison** | | | **Between group comparison** | | | |
| | | | | | | **group** | **Test** | **p** | **Test** | **p** | **Test** | **p** |
| **General knockdown** | | | | | | | | | | | | |
| active | BL, EXT | 1,18 | **group** | 0.002 | 0.97 | | | | | | **control** | |
| | | | **Test-session** | 33.75 | 0.0001 | **control** | BL, EXT | 0.003 | BL | 0.93 | BL | 0.002 |
| | | | **interaction** | 0.0077 | 0.93 | **KD** | BL, EXT | 0.0008 | EXT | 0.97 | EXT | 0.0002 |
| | EXT, RE | 1,18 | **group** | 0.007 | 0.94 | | | | | | | |
| | | | **Test-session** | 132.65 | 0.0001 | **control** | EXT, RE | 0.0002 | | | RE | 0.0001 |
| | | | **interaction** | 0.0003 | 0.99 | **KD** | EXT, RE | 0.0002 | RE | 0.97 | | |
| inactive | BL, EXT | 1,18 | **group** | 2.49 | 0.13 | | | | | | **KD** | |
| | | | **Test-session** | 14.73 | 0.001 | **control** | BL, EXT | 0.02 | BL | 0.3 | BL | 0.002 |
| | | | **interaction** | 0.19 | 0.67 | **KD** | BL, EXT | 0.07 | EXT | 0.13 | EXT | 0.009 |
| | EXT, RE | 1,18 | **group** | 3.99 | 0.06 | | | | | | | |
| | | | **Test-session** | 70.64 | 0.0001 | **control** | EXT, RE | 0.0002 | | | RE | 0.0001 |
| | | | **interaction** | 0.004 | 0.95 | **KD** | EXT, RE | 0.0002 | RE | 0.14 | | |
| **Cre-inducible knockdown** | | | | | | | | | | | | |
| active | BL, EXT | 1,12 | **group** | 0.272 | 0.611 | | | | | | **control** | |
| | | | **Test-session** | 23.77 | 0.0003 | **control** | BL, EXT | 0.0009 | BL | 0.58 | BL | 0.023 |
| | | | **interaction** | 0.005 | 0.82 | **KD** | BL, EXT | 0.017 | EXT | 0.81 | EXT | 0.02 |
| | EXT, RE | 1,12 | **group** | 0.089 | 0.77 | | | | | | | |
| | | | **Test-session** | 29.88 | 0.0001 | **control** | EXT, RE | 0.008 | | | RE | 0.008 |
| | | | **interaction** | 0.048 | 0.048 | **KD** | EXT, RE | 0.005 | RE | 0.92 | | |
| inactive | BL, EXT | 1,12 | **group** | 1.52 | 0.24 | | | | | | **KD** | |
| | | | **Test-session** | 1.19 | 0.3 | **control** | BL, EXT | 0.69 | BL | 0.73 | BL | 0.005 |
| | | | **interaction** | 0.28 | 0.61 | **KD** | BL, EXT | 0.28 | EXT | 0.39 | EXT | 0.49 |
| | EXT, RE | 1,12 | **group** | 1.01 | 0.34 | | | | | | | |
| | | | **Test-session** | 0.17 | 0.34 | **control** | EXT, RE | 0.22 | | | RE | 0.006 |
| | | | **interaction** | 2.03 | 0.18 | **KD** | EXT, RE | 0.45 | RE | 0.7 | | |

**Supplementary Table 2** Changes in body weight of animals during treatment with LY379268 calculated as difference in body weight before and after treatment days (%).

| LY379268 dose, mg/kg | Body weight change, % |
|---|---|
| 0 | +0.5 |
| 1 | +0.0 |
| 3 | +0.2 |

**Supplementary Table 3** Changes in body weight of animals during treatment with LY354740 calculated as difference in body weight before and after treatment days (%).

| LY354740 dose, mg/kg | Body weight change, % |
|---|---|
| 0 | -0.2 |
| 3 | +0.1 |
| 10 | +0.2 |

**Supplementary Table 4** Changes in body weight of animals during treatment with LY487379 calculated as difference in body weight before and after treatment days (%).

| LY487379 dose, mg/kg | Body weight change, % | |
|---|---|---|
| | Males | Females |
| 0 | +0.5 | +0.4 |
| 10 | +0.5 | |
| 30 | -0.5 | -1.9* |

* indicates significant differences from the vehicle group, $p < 0.05$.
